# Role of clathrin-mediated endocytosis in the use of heme and hemoglobin by the fungal pathogen *Cryptococcus neoformans*

**DOI:** 10.1101/352534

**Authors:** Gaurav Bairwa, Mélissa Caza, Linda Horianopoulos, Guanggan Hu, James Kronstad

## Abstract

Heme is a major source of iron for pathogens of humans, and its use is critical in determining the outcome of infection and disease. *Cryptococcus neoformans* is an encapsulated fungal pathogen that causes life-threatening infections in immunocompromised individuals. *C. neoformans* effectively uses heme as an iron source but the underlying mechanisms are poorly defined. Non-iron metalloporphyrins (MPPs) are toxic analogues of heme and are thought to enter microbial cells via endogenous heme acquisition systems. We therefore carried out a mutant screen for susceptibility against manganese metalloporphyrin (Mn MPP) to identify new components for heme uptake in *C. neoformans*. We identified several genes involved in signaling, DNA repair, sugar metabolism and trafficking that play important roles in susceptibility to Mn MPP and in the use of heme as an iron source. We focused on investigating the role of clathrin-mediated endocytosis (CME) and found that several components of CME including Chc1, Las17, Rvs161 and Rvs167 are required for growth on heme and hemoglobin, and for endocytosis and intracellular trafficking of these molecules. We show that the hemoglobin uptake process in *C. neoformans* involves clathrin heavy chain, Chc1, which appears to co-localize with hemoglobin containing-vesicles and to potentially assist in proper delivery of hemoglobin to the vacuole. Additionally, *C. neoformans* strains lacking Chc1, Las17, Rvs161, or Rvs167 were defective in the elaboration of several key virulence factors and a *las17* mutant was avirulent in a mouse model of cryptococcosis. Overall, this study unveils crucial functions of CME in the use of heme iron by *C. neoformans* and reveals a role for CME in fungal pathogenesis.

## Introduction

*Cryptococcus neoformans* is an encapsulated fungal pathogen that causes life-threatening infections in immunocompromised individuals, especially HIV/AIDS patients (Kronstad *et al.*, 2011; Kwon-Chung *et al.*, 2014). A recent study estimated that 223,100 individuals acquired cryptococcal meningitis globally in 2014, with a very high mortality rate of nearly 81% (Rajasingham *et al.*, 2017). The common virulence traits of *C. neoformans* include cell surface elaboration of capsule and melanin, production of extracellular enzymes such as urease and phospholipase B, and adaptation to the host environment including growth at 37°C, sensing nutritional cues and metal-ion availability, and evading the host-immune system (Djordjevic *et al.*, 2005; Doering, 2009; Djordjevic, 2010; Almeida *et al.*, 2015; Wager *et al.*, 2016; Campuzano and Wormley, 2018).

Sensing and adapting to the availability of metal ions such as iron and copper in the host environment are pivotal for any pathogen to survive and proliferate (Skaar, 2010; Hood and Skaar, 2012; Palmer and Skaar, 2016; Bairwa *et al.*, 2017). Iron is an essential metal required for several enzymatic and biochemical processes in eukaryotic cells, and the metal plays a crucial role in the virulence of microbial pathogens including *C. neoformans* (Jung and Kronstad, 2008; Jung *et al.*, 2008; Weinberg, 2009; Skaar, 2010; Bairwa *et al.*, 2017). The ability to use host iron sources, such as heme/hemoglobin and transferrin, is advantageous for the growth and survival of *C. neoformans* inside a vertebrate host (Jung *et al.*, 2008). Previous studies have shown key roles for the ESCRT proteins Vps23, Vps22, and Snf7 in the use of heme iron and in the virulence of *C. neoformans* (Hu *et al.*, 2013; da C. Godinho *et al.*, 2014; Hu *et al.*, 2015). Additionally, a putative hemophore, Cig1, was discovered to be involved in heme binding and uptake, and in virulence (Cadieux *et al.*, 2013). Despite some knowledge of the machinery for heme uptake and use in *C. neoformans*, the detailed mechanism of heme-iron uptake, trafficking, metabolism, storage, and regulation in this pathogen remain unknown.

Non-iron metalloporphyrins (MPPs) possess strong anti-bacterial, anti-fungal, and anti-parasitic activities (Stojiljkovic *et al.*, 1999; Stojiljkovic *et al.*, 2001). The uptake of these compounds in bacteria is thought to occur via high-affinity heme transport (Stojiljkovic *et al.*, 1999). Previous studies have shown that non-iron MPPs having metal ions such as gallium (Ga), manganese (Mn), zinc (Zn) ruthenium (Ru) or indium (In) at their coordination center, can have anti-bacterial activity against both gram-positive and gram-negative bacteria to varying degrees, with Ga-MPP as the most potent (Stojiljkovic *et al.*, 1999; Stojiljkovic *et al.*, 2001). Since the transport of non-iron MPPs occurs through heme uptake systems in bacteria, it may be possible to identify new components of the machinery for heme use in *C. neoformans* by characterizing mutants with altered susceptibility to non-iron MPPs.

Clathrin-mediated endocytosis (CME) is employed by eukaryotic cells to acquire receptor-bound cargo and proteins from the cell surface. A role for endocytosis in heme uptake has been previously described for the pathogenic yeast *Candida albicans* (Weissman *et al.*, 2008). Therefore, it is plausible that the CME may play a similar role in heme uptake in *C. neoformans*. The process of CME, beginning from cargo identification and coat assembly to coat scission and disassembly, is evolutionary conserved in eukaryotic cells with most of the proteins having well-characterized homologs in yeast and mammalian cells (Lu *et al.*, 2016; Kaksonen and Roux, 2018). However, some differences have been reported for CME in yeast compared to higher eukaryotes (Yeung *et al.*, 1999). CME involves the ordered temporal and spatial recruitment and organization of approximately 60 conserved proteins at the plasma membrane to internalize receptor-bound cargo in a clathrin-coated vesicle (CCV) (Kaksonen *et al.*, 2005; Mettlen and Danuser, 2014; Ferguson *et al.*, 2016). One of the first proteins to be recruited at the internalization site is clathrin and this protein is the polymeric assembly unit of a vesicular coat formed by three heavy chains and three light chains encoded by *CHC1* and CLC1 in yeast, respectively (Edeling *et al.*, 2006; Kirchhausen, 2009; McMahon and Boucrot, 2011). The clathrin assembly units have a triskelion shape and interact with each other to form a polyhedral lattice surrounding the vesicle. Clathrin adaptor proteins are involved in linking the clathrin coat with the phospholipids of vesicle membranes and/or cargo transmembrane proteins (Edeling *et al.*, 2006). The adaptor protein AP2 participates in the internalization of several cargo proteins in mammalian cells, including the well characterized transferrin receptor, but no such functions have been assigned to AP2 in yeast (Yeung *et al.*, 1999). Following the early stage of assembly of a clathrin coat at the cytoplasmic face of the plasma membrane, several additional proteins including Sla2, Ent1/Ent2 and Lsb3/4/5, Pan1, End3 and Sla1 are recruited at distinct times as the process moves into intermediate and late stages of coat assembly (Feliciano and Pietro, 2012; Mettlen and Danuser, 2014; Sun *et al.*, 2015). At the late stage of CCV formation, a group of proteins (Wiskott-Aldrich Syndrome Protein (WASP) and myosin) arrive to initiate CCV internalization (Tyler *et al.*, 2016). The Las17 protein, which is a yeast homolog of WASP in mammals, acts as a nucleation-promoting factor for actin assembly near the neck region to initiate the internalization of the CCV with Myosin-3/5 (Sun *et al.*, 2006; Robertson *et al.*, 2009). In the final stages of yeast endocytosis, scission of the vesicle from the plasma membrane is achieved by the BAR (Bin Amphiphysin Rvs) domain proteins, Rvs161 and Rvs167, along with the yeast dynamin-related protein Vps1 (Nannapaneni *et al.*, 2010; Rooij *et al.*, 2010; Rooij *et al.*, 2012). As with clathrin, other proteins associated with CCV formation are recycled into the cytosol to participate in subsequent rounds of clathrin-coated vesicle formation.

We report here the identification of several novel players in the machinery for heme use by *C. neoformans* through a screen for mutants with altered susceptibility to a non-iron metalloporphyrin. The screen identified genes for various processes, including vesicular trafficking, transcriptional regulation, and DNA repair and signaling, that are required for the robust use of heme iron by *C. neoformans*. In particular, we discovered a previously unidentified role of CME for heme use and found that multiple components of CME *viz*. Chc1, Las17, Rvs161 and Rvs167 are required for the endocytosis, intracellular trafficking, and use of hemin and hemoglobin. Importantly, we show that the clathrin heavy chain protein Chc1 is involved in internalization of hemoglobin and proper intracellular trafficking. Further characterization of the *C. neoformans* mutants lacking *CHC1* and *LAS17* revealed altered capsule and melanin production, and defects in several physiological growth conditions. Additionally, loss of Las17 resulted in attenuated virulence in a mouse inhalation model of cryptococcosis. Overall, this study provides important insights into the contributions of CME in heme use and more broadly in the pathogenesis of *C. neoformans*, and suggests the presence of a heme/hemoglobin receptor in this fungal pathogen.

## Results

### Identification of mutants with altered susceptibility to manganese metalloporphyrin

Non-iron metalloporphyrins (MPs) are toxic for microbes and their uptake is thought to occur via heme transport systems (Stojiljkovic *et al.*, 1999; Stojiljkovic *et al.*, 2001; Wakeman *et al.*, 2014; Mitra *et al.*, 2017). To establish conditions for screening mutants on MPs, we initially evaluated the susceptibility of a wild-type (WT) *C. neoformans* strain to two potent anti-bacterial MPs, manganese and gallium metalloporphyrin (MnMPP and GaMPP, respectively) (Fig. 1A). Additionally, we examined the growth of the WT strain with non-iron metalloporphyrins after iron starvation. For these experiments, we employed the minimal medium YNB, YNB with the iron chelator BPS (iron-starved condition), YNB+BPS supplemented with hemin (iron-rescued condition) and YNB+BPS supplemented with hemin and either MnMPP or GaMPP (metalloporphyrin condition). As shown in Figure 1A, the growth of non-starved WT cells in different concentrations of MnMPP or GaMPP was comparable to the condition with heme, suggesting that the concentrations evaluated were not toxic to WT cells containing sufficient iron. In contrast, prior starvation of WT cells for iron resulted in sensitivity to increasing concentrations of both MnMPP and GaMPP, with growth significantly reduced in the highest concentrations tested (Fig. 1A). We also observed that GaMPP was toxic for the iron-starved cells at lower concentrations compared to MnMPP. These results indicated that pre-starvation was necessary for mutant screening, and that MnMPP at 10 μM in the presence of hemin provided a condition to identify susceptible and (potentially) resistant mutants.

**Figure 1:**
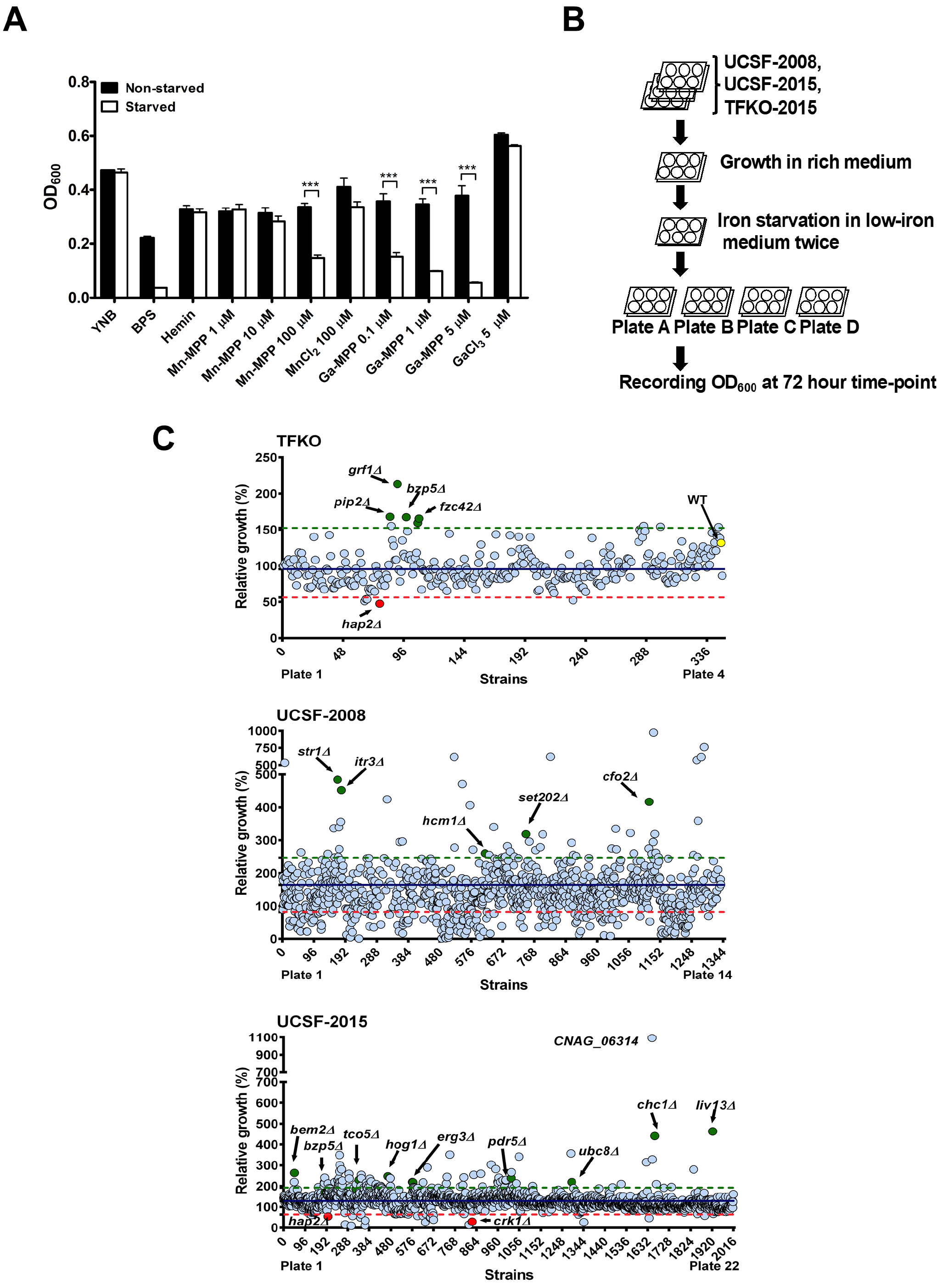
Identification of cryptococcal mutants with altered susceptibility to manganese metalloporphyrin (MnMPP). (A) Growth profile at 72 h for iron-starved or non-starved WT cells in YNB medium containing different concentrations of either manganese or gallium metalloporphyrin. The results are the means ± standard errors of the mean (SEM) from three independent experiments. (\*\*\**P* < 0.001). (B) Graphical outline of the strategy used to screen the deletion mutant libraries against MnMPP. Plate A; YNB, Plate B; YNB with BPS (150 μM), Plate C; YNB with BPS (150 μM) and hemin (10 μM) [hemin media] and Plate D; YNB with BPS (150 μM), hemin (10 μM) and MnMPP (10 μM) [MnMPP media]. (C) Growth profile of individual strains of the three libraries screened against MnMPP, representing the relative growth differences in percentage between the MnMPP and hemin media conditions at 72 h. The blue line represents the average relative growth (ARG) value calculated from the relative growth difference values of all the strains in a library. The red line marks the value which is 50% of the ARG, while the green line marks the value at 150% of the ARG. Circles filled with green color highlight some of the interesting mutants showing resistance to MnMPP. Red filled circles represent some of the interesting mutants that are sensitive to MnMPP.

We screened three independent libraries of *C. neoformans* strains to identify mutants with altered susceptibility to MnMPP. Two libraries of gene deletion mutants were generated by Dr. Hiten Madhani’s group at the University of California, San Francisco (designated the UCSF-2008 and UCSF-2015 collections, arranged in 96-well plates) and obtained from the Fungal Genetics Stock Center (FGSC, USA) (Liu *et al.*, 2008). A third collection designated the TFKO library consists of mutants deleted for 155 putative transcription factors of *C. neoformans* (Jung *et al.*, 2015). The two UCSF library collections of ~3,300 mutants were arranged in 36 plates and the transcription factor deletion strains were arranged in 4 plates. The strains were screened in a 96 well format using liquid YNB medium containing BPS and hemin, with and without MnMPP, as described in the Experimental Procedures. The screen was initiated by starving the mutants for iron for 2 days in YNB with BPS in the 96-well format, followed by inoculating the starved cells into test media and measuring their growth by absorbance at OD_600_ after 72 hours (Fig. 1B and described in Experimental Procedures). Figure 1C depicts the relative growth difference (in percentage) of individual strains (each denoted with a circle) in medium with hemin and MnMPP versus medium with only hemin for all three libraries. An average of relative growth difference values of all the strains in a library was determined. This average relative growth (ARG) value is denoted by the solid blue line, while the green and red dotted lines represent the cut-off for relative growth difference values that are 150% higher or 50% lower than the ARG. The strains that grew ≥ 150% more than the ARG were selected as resistant candidates, while strains growing ≤ 50% of the ARG values were selected as MnMPP sensitive mutants (Fig. 1C and Supporting Information Table S1). Overall, the screens yielded both resistant and susceptible mutants, and we focused on the strains showing resistant to MnMPP toxicity for subsequent analysis. Our rationale was that these mutants might identify functions necessary for the uptake or use of hemin, as well as targets of MnMPP toxicity.

As shown in Figure 1C, four mutants were identified in the TFKO library as resistant strains (*pip2*Δ, *grf1*Δ, *bzp5*Δ and *fzc42*Δ) and *hap2*Δ was identified as a sensitive mutant. Pip2 (*CNAG_05170*) is a fungal specific Zn(2)-Cys(6) binuclear cluster domain transcription factor and Grf1 (*CNAG_05186*) is a GRF zinc finger protein. Bzp5 (*CNAG_07940*) is a basic leucine zipper transcription factor and Fzc42 (*CNAG_05112*) is a hypothetical protein with a Zn(2)-Cys(6) domain (Jung *et al.*, 2015). Hap2 (*CNAG_07435*) is a CCAAT-binding transcription factor and is an essential component of Hap2/Hap3/Hap4/Hap5 CCAAT-binding complex involved in global regulation of respiratory gene expression in *Saccharomyces cerevisiae* (Olesen and Guarente, 1990). In the fungal pathogen *C. albicans*, Hap2 has been found to be involved in regulation of low-iron induction of the *FRP1* gene which encodes a ferric reductase (Baek *et al.*, 2008). Hap2 in *C. neoformans* is thought to interact with the Hap3, Hap5 and HapX proteins that were previously shown to be required for robust growth on heme (Jung *et al.*, 2010).

The screens for the two UCSF libraries yielded a large number of mutants that were either sensitive or resistant to MnMPP, and a number of these are highlighted in Figure 1C. We did detect common mutants that were present in all three libraries in support of the fidelity of the screening approach. For example, we found the *bzp5*Δ and *hap2*Δ mutants in both the TFKO and UCSF-2015 libraries. Furthermore, two strains deleted for the genes *CNAG_06601* or *CNAG_00264* were identified as resistant in both the UCSF-2008 and UCSF-2015 libraries. Other than these examples, the candidate mutants that we identified were exclusive to one of the libraries. Notably the screens identified factors already known to be involved in iron use by *C. neoformans*. These included mutants for the genes *STR1* (encoding a putative siderophore transporter), *CFO2* (encoding a ferroxidase) and as well as *HAP2* mentioned above (Jung *et al.*, 2008; Jung *et al.*, 2009; Jung *et al.*, 2010). These results support the idea that the screens captured factors relevant to iron acquisition and homeostasis.

### Mutants with resistance to MnMPP represent diverse GO categories and are enriched for functions in sugar metabolism, DNA synthesis and repair, and transcriptional regulation

We concentrated on the resistant collection of mutants in our subsequent analyses in this study. In total, 151 mutants were obtained from the two FGSC libraries and 4 mutants were identified from the TFKO library; these were analyzed for their functional annotations and potential interactions. Of the 155 mutants, two corresponding genes, *CNAG_03546* and *CNAG_03505*, could not be identified in the FungiDB database (http://fungidb.org). To perform Gene Ontology (GO) and *STRING* analyses, we retrieved the orthologous *C. neoformans* JEC21 gene names for 146 of the remaining 153 genes from the FungiDB database. Unlike the serotype A strain of *C. neoformans* that was used for our screen, the JEC21 genome information has been organized to support GO and *STRING* analyses, and the JEC21 gene designations were used for the presentation of the data in Figure 2C.

The GO analysis revealed a large representation of the genes involved in metabolic processes (31%), response to stimulus (10%) and biological regulation (5%). Among these, there was enrichment for genes involved in glucose metabolism and especially gluconeogenesis (e.g., *CNB04050* encoding glucose-6-phosphate isomerase and *CNF00650* encoding pyruvate carboxylase) (Figs. 2A and B). Furthermore, a large number of genes in the category of response to stimulus were involved in the cellular response to DNA damage and mismatch repair (e.g., *FEN1* encoding Flap endonuclease, *MSH2/CNA07480*, *CNA03100* encoding a single-stranded DNA specific endodeoxyribonuclease, and two uncharacterized genes: *CNK01700* and *CND04530*), suggesting that DNA stress is a substantial component of MnMPP toxicity. One interesting category of genes was involved in clathrin-vesicle coat and the associated processes of endocytosis and trafficking. This group was enriched in the list of mutants conferring MnMPP resistance and included *CNJ03270* encoding the clathrin heavy chain, *CNA03420* encoding Hob1 and *CNE02750* encoding the autophagy protein, Atg18 (Fig. 2B).

A *STRING* network analysis was performed to reveal clusters of interactions between functionally associated gene products. Most notable interactions and clustering were obtained for the genes involved in: 1) Hog1-related stress signaling including *HOG1* which encodes a mitogen-activated protein (MAP) kinase, *CNB1* encoding the calcineurin subunit B, *CNA07470/PBS2* encoding a MAP kinase kinase, *SCH9* encoding a protein kinase and *CNH01060* encoding a calmodulin-dependent protein kinase-I; 2) Functions for gluconeogenesis (e.g., *CNA00280* encoding an alcohol dehydrogenase, *CNF03900* encoding an aldehyde dehydrogenase, *CND03740* encoding aldo-keto reductase) and; 3) DNA synthesis and mismatch repair (e.g., *FEN1*, *MSH2*, *CNA03100*, *CNG03940*, *CNK01700* and *CND04530)* (Fig. 2C). Another small cluster contained genes encoding candidate transporters including *CNL06490* (*ABC* transporter), *CNC00270* (a multi-drug resistance protein), and *CNC03290* (tetracycline efflux protein) (Fig. 2C). Taken together, these results illustrate the diversity of functions targeted by MnMPPs and suggest that the compound provokes stress and metabolic responses.

**Figure 2:**
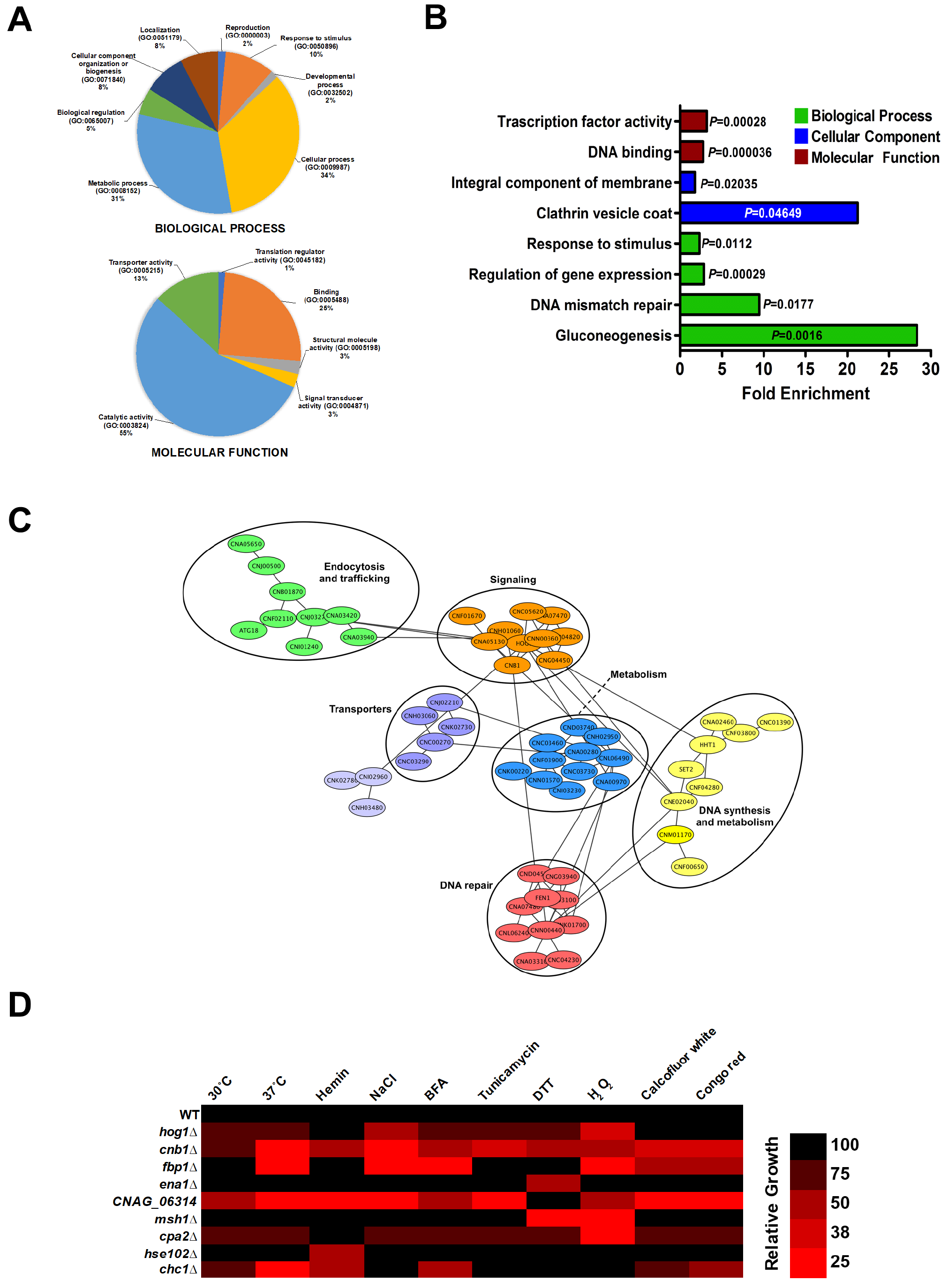
Gene Ontology and interaction analyses of genes involved in MnMPP resistance. (A) Gene Ontology analysis of the genes involved in MnMPP resistance for the categories of biological process and molecular functions. The H99 genes were matched to the *C. neoformans* JEC21 gene IDs retrieved from the FungiDB (http://fungidb.org/fungidb/) database and analyzed with the PANTHER gene ontology online tool (http://pantherdb.org) using *C. neoformans* as the organism. (B) Functional enrichment analysis of genes involved in MnMPP resistance. Genes were analyzed for enrichment of the GO biological process, molecular function and cellular component categories. Enriched groups were scored by comparison to a background list of the *C. neoformans* H99 genome database using a cut-off of *P* < 0.05 and Bonferroni correction. (C) STRING network analysis of the *C. neoformans* genes involved in MnMPP resistance. Interaction network details were obtained with JEC21 gene IDs from the STRING online tool (https://string-db.org). The network was extracted and visualized with Cytoscape 3.6.1 software (http://www.cytoscape.org) and clustering was performed with the clusterMaper plugin using the community cluster (GLay) network algorithm. Individual clusters are marked with an elliptical circle. (D) Heatmap depicting the growth behavior of selected MnMPP resistant mutants in various environmental stress conditions. Growth of the individual strains at 30°C, 37°C and in presence of hemin (10 μM), NaCl (1.5 M), brefeldin A; BFA (20 μg ml^−1^), tunicamycin (0.15 μg ml^−1^), dithiothreitol; DTT (10 mM), H_2_O_2_ (1 mM), calcofluor white (1 mg ml^−1^), and congo red (2.5 mg ml^−1^) was checked and displayed with color gradients (indicated as percentages of the WT).

### MnMPP resistant mutants show sensitivity to a diverse set of stress conditions

To better understand the functional importance of the genes involved in the resistance to MnMPP, we characterized 50 MnMPP resistant mutants for growth under diverse environmental conditions including host temperature (37°C), high salt (e.g., NaCl), trafficking inhibition (brefeldin A), cell wall damage (calcofluor white and congo red), challenges to protein folding (tunicamycin and dithiothreitol) and stress from reactive oxygen species (H_2_O_2_). As shown in Fig. 2D and Supporting Information Fig. S1, several of the mutants (e.g., *hog1*Δ, *cnb1*Δ, *cpa2*Δ, *ade5*Δ, and strains deleted for *CNAG_04091, CNAG_05159* and *CNAG_06314*) showed susceptibility to a variety of stress conditions. For the strains lacking *HOG1* or *CNB1*, the defect at 37°C has been previously observed (Fox *et al.*, 2001; Bahn *et al.*, 2005). Among the 50 mutants tested, 10 strains showed susceptibility to at least five stress conditions (Supporting Information Fig. S1). Importantly, we did not observe any common phenotypic themes for the strains deleted in the same functional classes of genes. That is, the phenotypic analysis of the selected strains did not reveal enrichment for any specific growth defects under any of the conditions tested.

### Chc1 is required to use heme-iron sources in C. neoformans

Our screen revealed an important role for genes encoding endocytic and trafficking functions in the susceptibility of *C. neoformans* to MnMPP. Given that the uptake of non-iron metalloporphyrins is thought to require heme transport machinery in bacteria (Stojiljkovic *et al.*, 1999), we carried out a detailed phenotypic characterization of *CHC1* (*CNAG_04904*) which encodes the clathrin heavy chain with a well-established role in endocytosis. *CHC1* is an evolutionary conserved gene and is not essential in *C. neoformans*, unlike the situation in higher eukaryotes. *CHC1* in *C. neoformans* encodes a 1,684 amino acid protein containing seven <u>C</u>lathrin <u>H</u>eavy <u>C</u>hain <u>R</u>epeat (CHCR) sequences of 145-amino acid residues each (Fig. 3A). These repeats are arranged tandemly in the C terminal portion of the protein and are evolutionary conserved in different kingdoms (Fig. 3A). Earlier studies have shown that the CHCR repeat is involved in protein-protein interactions or clathrin binding (Ybe *et al.*, 1999; Young, 2007). Notably, our phylogenetic analysis revealed greater similarity between Chc1 of Cryptococcus *spp*. and mammalian counterparts compared to the Chc1 proteins of the yeasts *S. cerevisiae* and *C. albicans* (Fig. 3B). As reported for *S. cerevisiae*, deletion of *CHC1* in *C. neoformans* also resulted in abnormal cell morphology, defects in cell division, and chitin deposition at the mother-daughter bud neck (Fig. 4A). Our phenotypic analysis with the *chc1*Δ mutant also showed that its growth was inhibited by the antifungal drug fluconazole (Fig. 4B). Furthermore, supplementation with hemin in the fluconazole medium did not rescue the growth defect of the *chc1*Δ mutant. In contrast, growth of the WT strain could be rescued by hemin even at higher fluconazole (10 μg ml^−1^) concentrations (Fig. 4B). This result suggested that the *chc1*Δ mutant was unable to efficiently take up hemin to rescue the activity of the heme-dependent lanosterol demethylase targeted by fluconazole.

**Figure 3:**
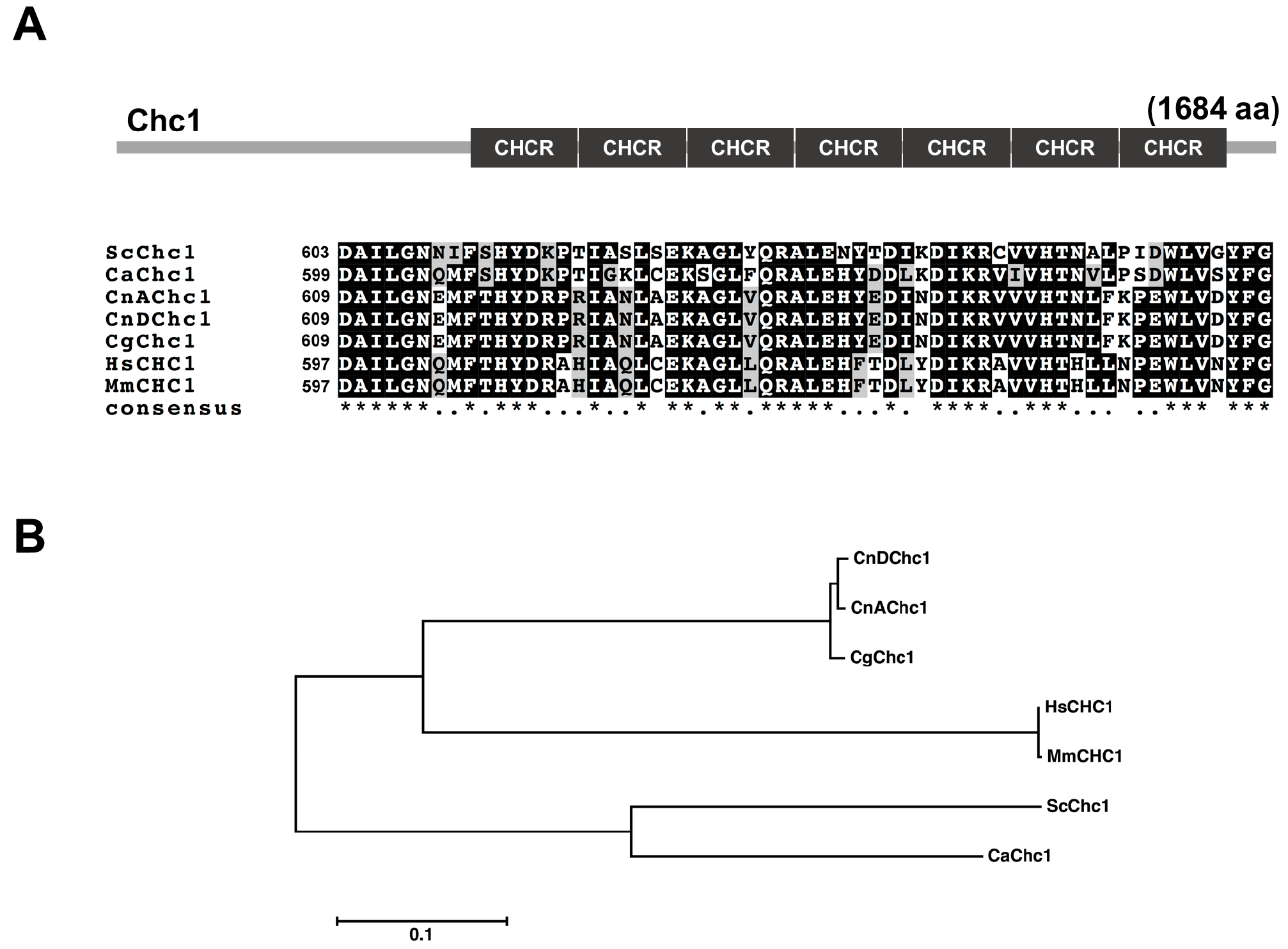
Sequence analysis of the clathrin heavy chain 1 protein in fungi and higher eukaryotes. (A) Predicted domain architecture of the *C. neoformans* Chc1 protein and multiple alignment of the 1^st^ Clathrin Heavy Chain Repeat (CHCR) sequences of the Chc1 proteins from selected organisms. The amino acid sequence alignment was performed with Clustal W (http://www.ch.embnet.org); identical residues are boxed in black (indicated with *) and similar residues are shown in gray (indicated with •). The numbers on the left represent the starting position of the CHCR repeat sequence in the protein from the respective organism. (B) Phylogenetic tree of the Chc1 proteins from selected organisms. The phylogenetic analysis of the Chc1 protein was performed in MEGA7 (https://www.megasoftware.net) by using the Maximum Likelihood method and applying Neighbor-Join and BioNJ algorithms. The tree is drawn to scale, with branch lengths measured in the number of substitution per site. *CNA, C. neoformans* var. *grubii, H99 (serotype A); CnD, C. neoformans* var. *neoformans JEC21 (serotype D); Cg, C. gattii; Sc, Saccharomyces cerevisiae; Ca, Candida albicans; Hm, Homo sapiens; Mm, Mus musculus.*

The possible role for Chc1 in hemin uptake prompted us to evaluate the role of the protein in iron acquisition by analyzing the growth of the *chc1*Δ strain in iron-starved conditions and in the presence of different iron sources. We first generated two independent *chc1*Δ (*chc1*Δ-*1* and *chc1*Δ-*2*) mutants in the *C. neoformans* H99 (serotype A) background using homologous recombination. Next, we starved the WT, *chc1*Δ and complemented strains for iron by growing them in YNB medium with BPS (YNB-LIM). We then tested growth by spot assays on YNB and YNB-LIM agar plates with and without inorganic iron sources (FeCl_3_ or FeSO_4_) or hemin. As shown in Figure 4C, the iron-starved WT, *chc1*Δ and the complemented strains could grow well on rich medium (YPD) and the minimal medium YNB, but not on medium depleted for iron (YNB-LIM). Supplementing YNB-LIM with FeCl_3_, FeSO_4_ or hemin allowed the WT and complemented strains to grow similar to the levels seen on YPD and YNB. However, the *chc1*Δ mutants could only grow on YNB-LIM supplemented with either FeCl_3_ or FeSO_4_ but not with hemin at any concentration. We also performed the growth assays in liquid YNB media supplemented with hemin or hemoglobin. As shown in Figure 5, the *chc1*Δ mutants could not grow to the level of the WT strain on any of the concentrations of hemin and hemoglobin tested. Reintroduction of *CHC1* in *chc1*Δ strain completely rescued the growth defects on the low-iron media supplemented with heme or hemoglobin (Figs. 4C and 5). Additionally, we found that the *chc1*Δ mutants showed increased sensitivity to the drug curcumin, an iron-chelator found in turmeric, a traditional Indian spice and medicine (Supporting Information Fig. S2). Importantly, addition of the inorganic iron compounds FeCl_3_ and FeSO_4_ but not hemin could rescue the growth defect of the *chc1*Δ mutants on curcumin. All together, these findings support a role for Chc1 in the use of heme iron by *C. neoformans*.

**Figure 4:**
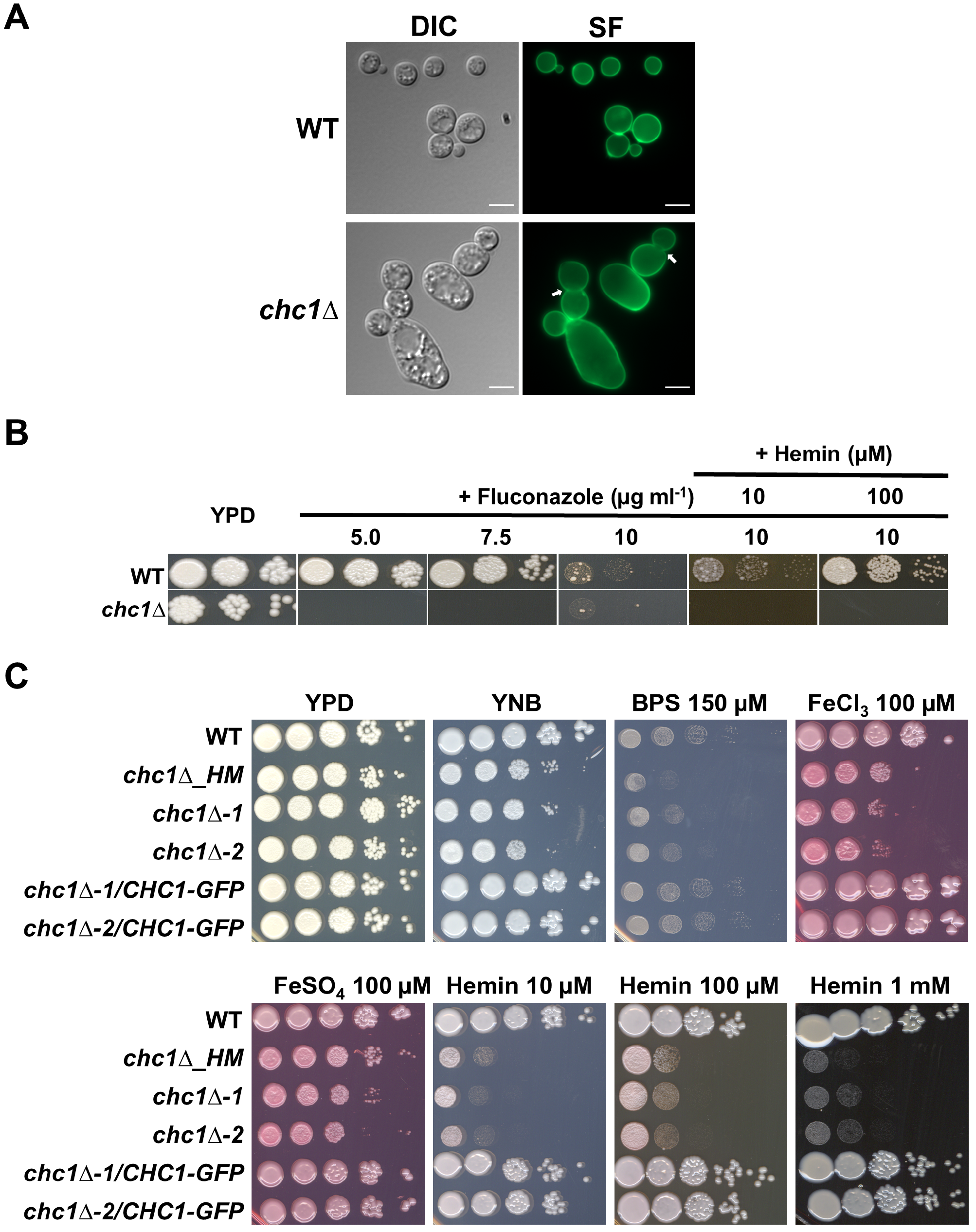
A *C. neoformans chc1*Δ mutant exhibits altered cell morphology and an inability to grow in presence of fluconazole and heme-iron sources. (A) Representative differential interference contrast (DIC) and fluorescent images of WT and *chc1*Δ cells stained with Solophenyl Flavine (SF) to visualize the cell architecture and chitin at the cell wall. White arrows indicate absence of chitin deposition at the mother-daughter bud neck of *chc1*Δ mutant strain. Scale bar = 5 μm. (B) and (C) Spot assays of 10-fold serial dilutions of the indicated strains on medium supplemented as shown. Cells of the indicated strains were starved for iron for 24 h prior to the spotting. All the plates were incubated for 3-4 days at 30°C before being photographed. The mutant designated *chc1*Δ_*HM* is from the UCSF-2015 deletion collection and the other two *chc1*Δ mutants were constructed as described in the Experimental Procedures. YNB; Yeast Nitrogen Base, BPS; Bathophenanthroline disulfonate.

**Figure 5:**
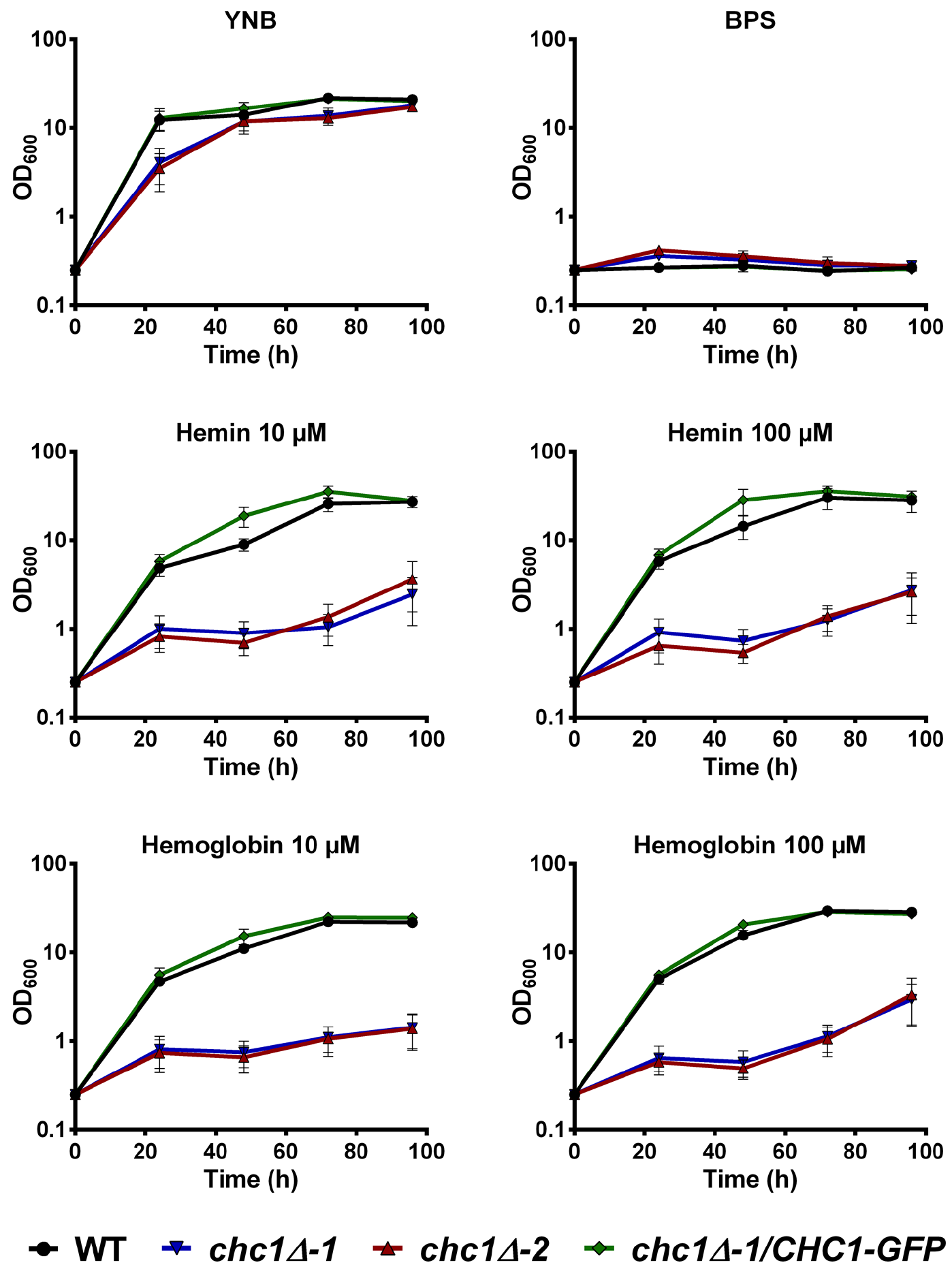
The *C. neoformans chc1*Δ mutant cannot use hemin and hemoglobin as sole iron sources. Cells of the indicated strains were starved for iron for 24 h and then transferred to YNB-LIM medium containing either hemin or hemoglobin. Growth at 30°C was monitored by recording OD_600_ over a 96-h time course at the indicated time intervals. Data are represented as the mean ± SEM of 3-4 independent experiments.

### Loss of Chc1 impairs endocytosis and trafficking

We next examined the role of Chc1 in endocytosis and trafficking in *C. neoformans*. In *S. cerevisiae*, Chc1 is a key component of clathrin-mediated endocytosis and plays an important role in internalization of several factors such as receptor-bound α-factor involved in mating and in the proper localization of several proteins (Baggett and Wendland, 2001). We initiated a characterization of Chc1 by tagging the protein at the C-terminus with GFP to assess its location in cells grown in YNB and low-iron YNB with and without hemin. The strain expressing Chc1-GFP had no discernable phenotypic defects in comparison to the WT strain. Importantly, introduction of the GFP-tagged version of Chc1 in the *chc1*Δ mutant fully rescued the growth defects of the strain (Figs. 4C and 5). As shown in Figure 6A, the Chc1-GFP protein was localized in punctate structures in cells grown in YNB and we hypothesize that these structures represent the trans-Golgi network (TGN), as has been previously shown for Chc1 in *S. cerevisiae*. Clathrin is known to mediate cargo selection and vesicle fission at the TGN and treatment with brefeldin A, an inhibitor of endoplasmic reticulum to Golgi trafficking, impairs assembly of the clathrin-dependent Golgi-apparatus (Radulescu *et al.*, 2007). Therefore, we treated the cells expressing Chc1-GFP with BFA to determine whether localization of Chc1-GFP was altered. Incubation of the cells with BFA for an hour caused reorganization of the Chc1-GFP puncta from the cytosol to the plasma membrane (Supporting Information Fig. S3A). That is, nearly 40% of the cells showed an increase in plasma membrane localization of Chc1-GFP after treatment with BFA (Supporting Information Fig. S3B). This result suggests that Chc1-GFP in *C. neoformans* localizes to the TGN or endocytic vesicles similar to Chc1 in *S. cerevisiae*. We also examined whether localization was altered by incubating cells in YNB-LIM with or without hemin. Our time course analysis did not reveal any differences in the localization of Chc1-GFP under the conditions examined (Supporting Information Fig. S3C).

**Figure 6:**
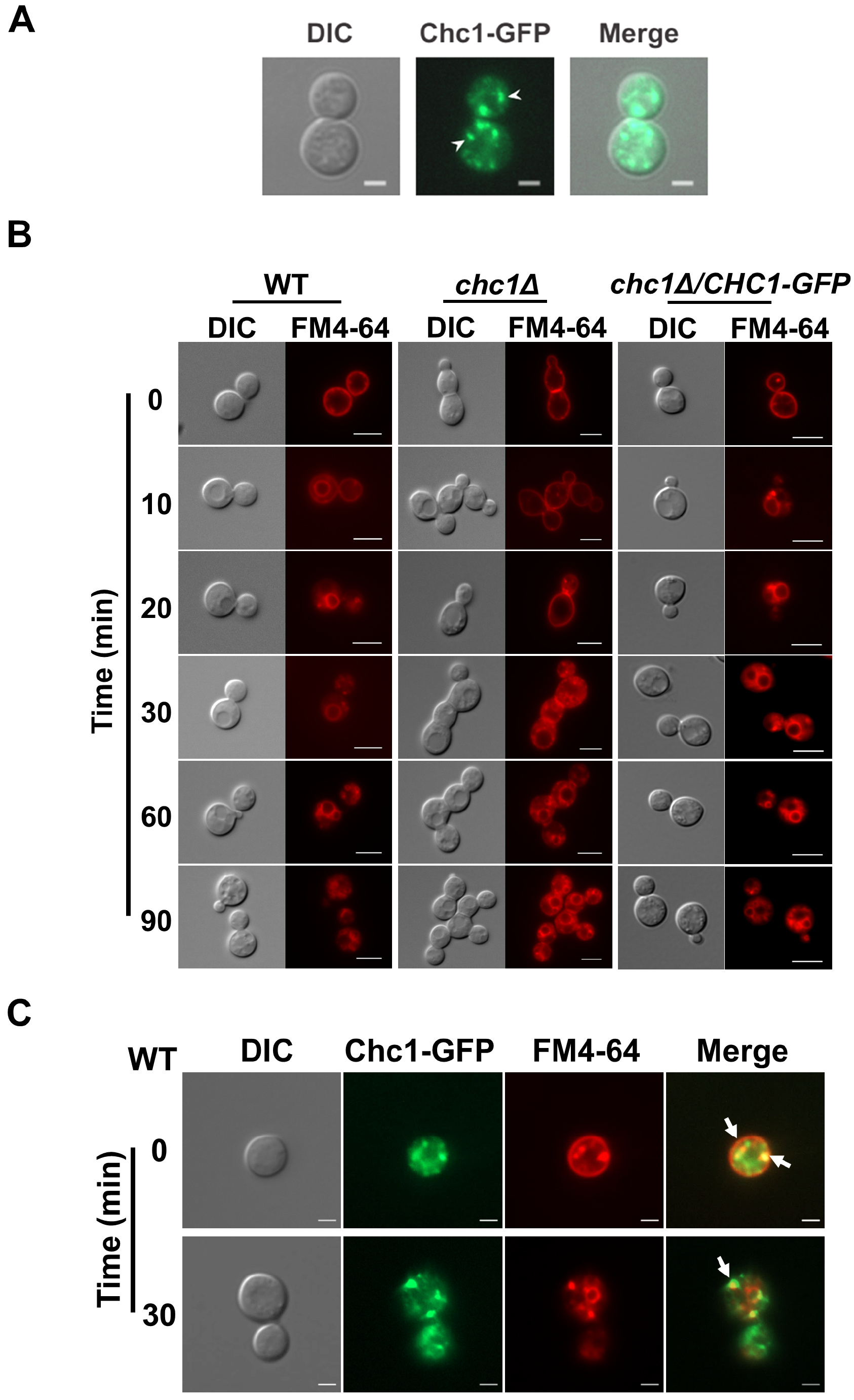
The *C. neoformans* Chc1 protein is required for endocytosis and intracellular trafficking. (A) *C. neoformans* Chc1 was tagged with GFP at its C-terminus and its localization was visualized in exponentially growing cells under a fluorescence microscope using the EGFP filter. White arrowheads indicate the Chc1-GFP puncta. Images are the representative of two independent experiments. Scale bar = 2 μm. (B) The absence of *CHC1* results in a significant delay in FM4-64 endocytosis and trafficking to the vacuole. YPD-grown cells of the indicated strains were labelled with FM4-64 on ice for 20 min and were visualized under a fluorescence microscope after the indicated times of incubation at room temperature. FM4-64 was visualized using a rhodamine filter. Images are representative of two independent experiments. Scale bar = 2 μm. (C) Chc1 in *C. neoformans* colocalizes with FM4-64-labelled structures. The WT strain expressing Chc1-GFP was labelled with FM4-64 and visualized under a fluorescence microscope after the indicated times of incubation at room temperature. Images are the representative of two independent experiments. Scale bar = 2 μm.

We further examined the role of Chc1 in endocytosis by performing assays with FM4-64, a lipophilic styryl dye which binds to the plasma membrane and is then endocytosed and trafficked to the vacuolar membrane (Vida and Emr, 1995). A defect in endocytosis either prevents or delays the trafficking of FM4-64 to the vacuole membrane. As shown in Figure 6B, most of the FM4-64 dye was internalized within 10 min of binding to the plasma membrane in WT cells. Furthermore, trafficking of internalized FM4-64 dye to the vacuolar membrane was completed by 30 min in the WT cells. In contrast, the *chc1*Δ mutant showed a significant delay in FM4-64 endocytosis and trafficking. Specifically, the internalization of plasma membrane-bound FM4-64 took more than 20 min, and trafficking to the vacuolar membrane was only partially completed by 90 min, a rate that was significantly (~3-fold) slower than in WT cells. Importantly, the defect in endocytosis of FM4-64 in the *chc1*Δ mutant was completely recovered to the WT level after complementation with the *CHC1-GFP* gene fusion at its native locus. To further examine the dynamics of Chc1-GFP localization during endocytosis, we followed the FM4-64 trafficking in the cells expressing Chc1-GFP. We observed that a portion of the Chc1-GFP puncta showed co-localization with FM4-64-labelled structures at both 0 and 30 min of time-course analysis (Fig. 6C). Taken together, these data indicate that Chc1 in *C. neoformans* plays an important role in endocytosis, as described for the protein in other eukaryotes.

### Chc1 is required for the uptake and trafficking of hemoglobin

Hemoglobin and heme-containing proteins provide a major potential source of iron for pathogens because ~ 80% of iron resides in heme in vertebrate hosts. Pathogens have therefore developed mechanisms to obtain iron from heme and hemoglobin to support proliferation during infection (Hood and Skaar, 2012; Palmer and Skaar, 2016). We previously demonstrated that *C. neoformans* can use heme and hemoglobin as sources of iron (Jung *et al.*, 2008). We hypothesized that Chc1 would affect the uptake or endocytosis of hemoglobin given that mutants lacking Chc1 failed to robustly proliferate with this iron source. To examine the use of hemoglobin in more detail, we labelled the human protein with the fluorescent dye Alexa Fluor 594 so that we could examine the kinetics of uptake and internalization by *C. neoformans* cells. Iron-starved cells from the WT, *chc1*Δ and complemented strains were incubated in YNB-LIM containing Alexa Fluor 594-labelled hemoglobin (Hb^Alexa594^) for 20 min on ice. The cells were then shifted to room temperature and observed under a fluorescence microscope to visualize the Hb^Alexa594^ cell-surface binding, uptake and trafficking at different time intervals. As shown in Figure 7A, labelled Hb^Alexa594^ could be observed on the cell surface of the WT, *chc1*Δ and *chc1*Δ/*CHC1-GFP* complemented strains after 5 minutes. By 30 min, we observed that most of the Hb^Alexa594^ was internalized and endocytosed into several small endocytic vesicles in both the WT and *chc1*Δ/*CHC1-GFP* complemented strains. In contrast, internalization of the cell surface bound Hb^Alexa594^ was significantly impaired in the *chc1*Δ and most of the labeled hemoglobin appeared to remain on the cell surface, even after 60 min.

To better understand the role of Chc1 in hemoglobin uptake, we also followed the internalization of Hb^Alexa594^ in the WT cells expressing Chc1-GFP. As shown in Figure 7B, we observed some co-localization of Chc1-GFP and Hb^Alexa594^ puncta after 30 min of labelling, but it was notable that not all of the Chc1-GFP puncta were co-localized with the Hb^Alexa594^-containing vesicles. Thus, there may be additional trafficking pathways for hemoglobin in *C. neoformans* including the involvement of ESCRT complexes (Hu *et al.*, 2015). We also hypothesized that the poor ability of the *chc1*Δ mutant to proliferate in different heme-iron sources and the gross defect in internalization of hemoglobin would result in reduced heme levels in the *chc1*Δ cells. As expected, the total heme content of the cells in *chc1*Δ strain was reduced 2-fold compared to the WT strain, and the heme levels were restored back to the WT levels in the *chc1*Δ/*CHC1-GFP* complemented strain (Fig. 7C). Overall these results highlight a contribution of Chc1 in hemoglobin uptake, endocytosis and trafficking.

**Figure 7:**
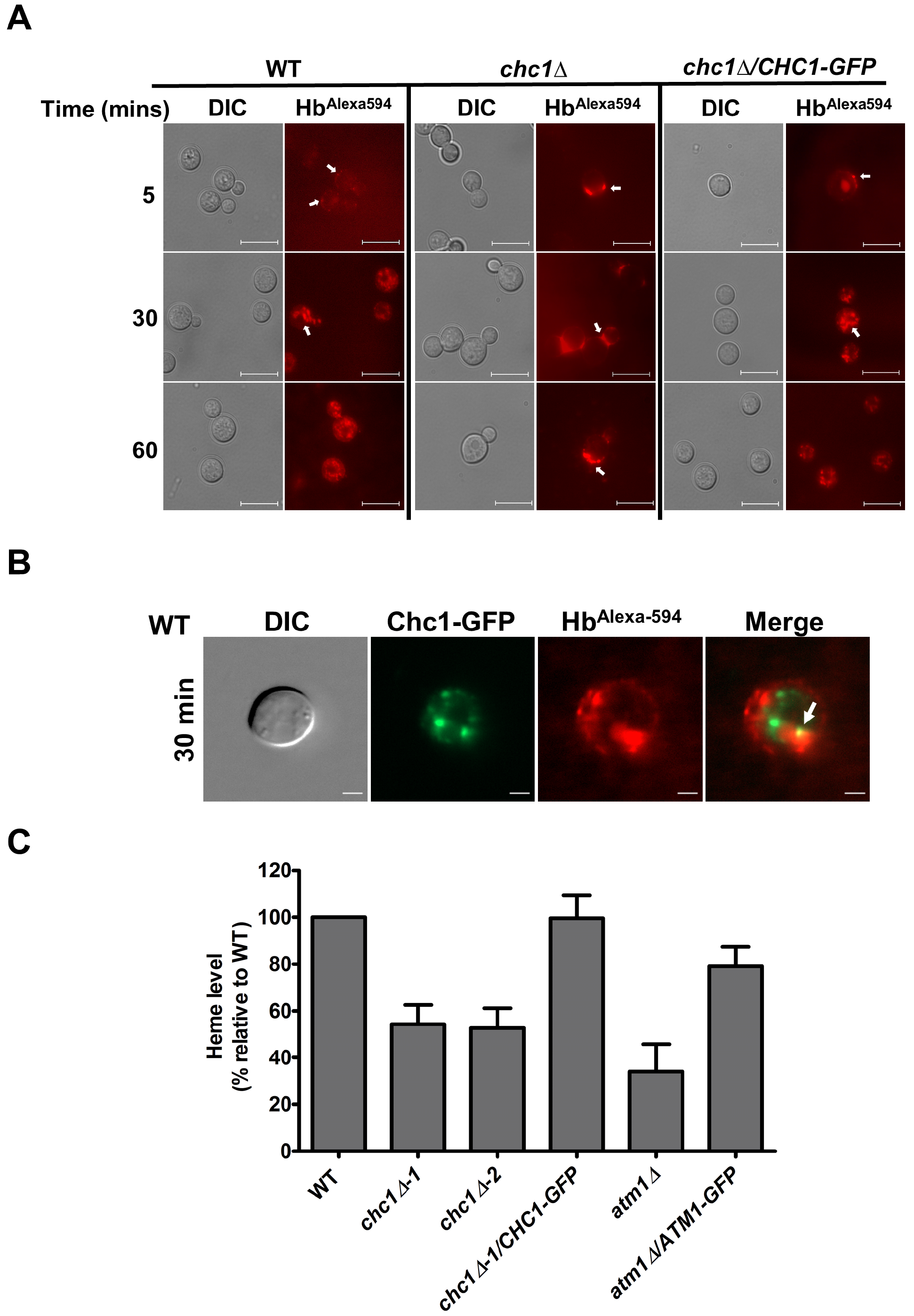
Chc1 is required for the endocytosis and trafficking of hemoglobin in *C. neoformans*. (A) To assess the uptake of labelled hemoglobin (Hb^Alexa594^), WT, *chc1*Δ and *chc1*Δ/*CHC1-GFP* complemented strains were incubated with Hb^Alexa594^ at 4°C and subsequently shifted to room temperature for the indicated times. Representative images of three independent experiments are shown. White arrows indicate the presence of Hb^Alexa594^. Scale bar = 5 μm. (B) Chc1 in *C. neoformans* colocalizes with hemoglobin containing vesicles. The WT strain expressing Chc1-GFP was incubated with Hb^Alexa594^ and visualized under a fluorescence microscope after 30 min incubation at room temperature. Representative images of two independent experiments are shown. The white arrow indicates the colocalization of Chc1-GFP and Hb^Alexa594^. Scale bar = 2 μm. (C) The *chc1* Δ strain exhibits reduced intracellular heme levels. Total heme content was measured and represented relative to the WT levels. *C. neoformans atm1*Δ and *atm1*Δ/*atm1-GFP* strains were used as the controls. Data represent the mean ± SEM of 4 independent experiments.

### Loss of Chc1 affects the elaboration of key virulence factors

*C. neoformans* survival inside the human host depends on elaboration of several key virulence determinants such as capsule and melanin production and the ability to grow at 37°C. We examined these phenotypes and found that deletion of *CHC1* completely abolished growth at 37°C. We characterized the *chc1*Δ mutant for capsule elaboration by growing cells in capsule inducing medium and observing capsule size by negative staining with India ink. Despite having a comparatively larger cell size, the capsule size of the *chc1*Δ mutant was reduced by 2-fold compared to the WT strain (Figs. 8A and B). Additionally, melanin production was reduced in the *chc1*Δ mutant compared to the WT strain on both solid and liquid L-DOPA media (Figs. 8C and D). Importantly, all these defects for the *chc1*Δ mutant were recovered to the WT level in the *chc1*Δ/*CHC1-GFP* complemented strain.

To further understand the role of Chc1, we analyzed the phenotypes of the *chc1*Δ mutant under different stress conditions including salt, osmotic and oxidative stresses, inhibition of secretion, use of alternate carbon sources, stressors of the endoplasmic reticulum/protein folding, and agents that challenged cell wall integrity. As shown in Figure 8E, lack of *CHC1* caused significant growth inhibition on the salts NaCl, KCl and LiCl. Sensitivity to LiCl was consistent with the reduced capsule phenotype of the *chc1*Δ mutant because we previously found a connection between capsule and LiCl (Hu *et al.*, 2007). Furthermore, the *chc1*Δ strain was unable to grow in presence of the secretion inhibitors brefeldin A (BFA) and monensin, and this is consistent with a role for Chc1 in intracellular trafficking. We also observed significant growth inhibition of the *chc1*Δ mutant in the presence of calcofluor white (CFW) and congo red that challenge cell wall integrity. Additionally, the *chc1*Δ mutant grew poorly in the presence of SDS, suggesting an altered plasma membrane composition (data not shown). Notably, growth of the *chc1*Δ mutant was similar to the WT strain on alternate carbon sources, the ER stress agents tunicamycin and DTT, different pH conditions and in response to oxidative/nitrosative stresses. Taken together these results suggest that in addition to the role in endocytosis and intracellular trafficking, Chc1 plays important role in the response to salt, osmotic and temperature stress, and contributes to intracellular trafficking and cell wall physiology in *C. neoformans*.

**Figure 8:**
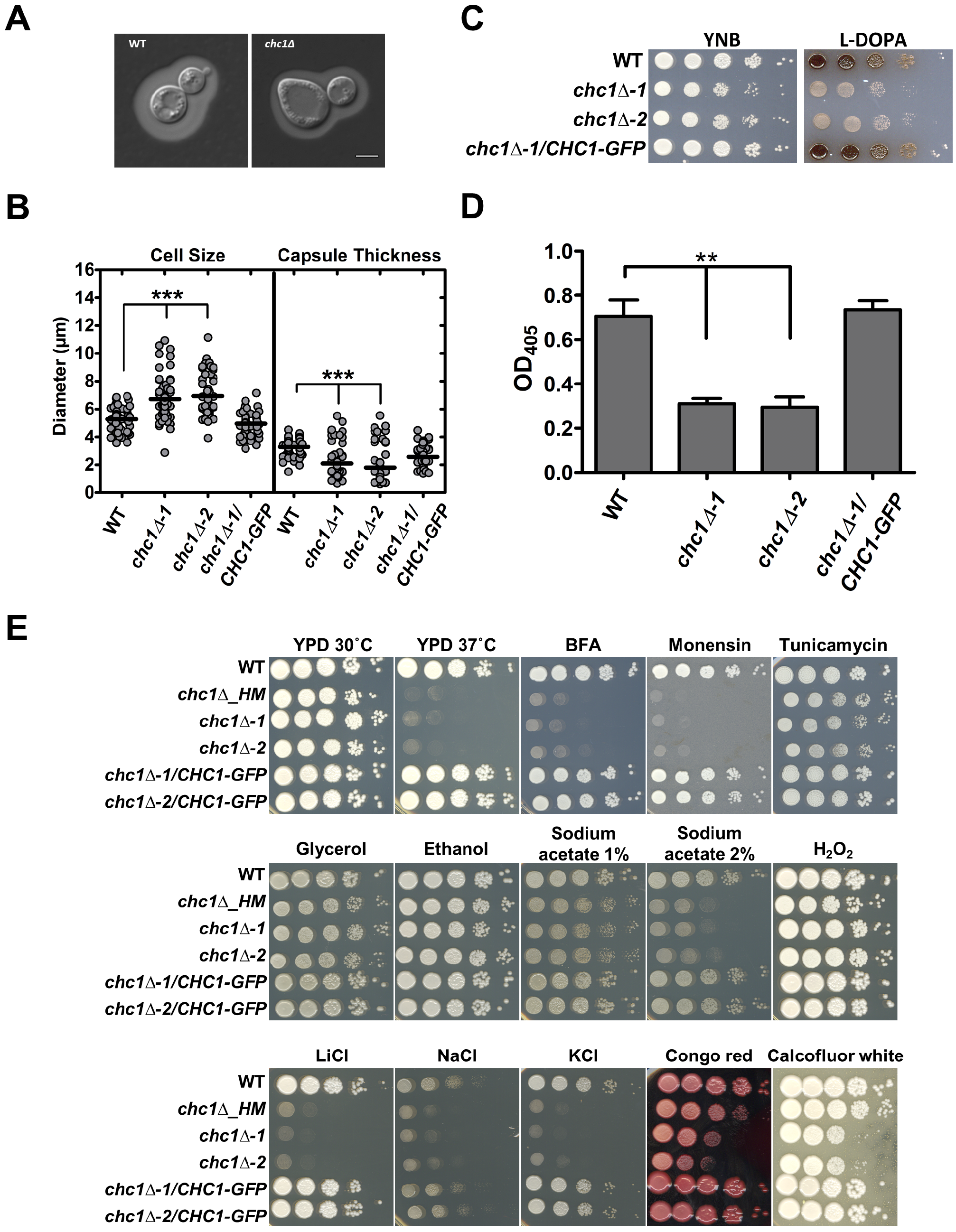
The *C. neoformans chc1*Δ mutant exhibits diminished capsule and melanin production and is sensitive to several stress conditions. (A) DIC microscopy images of India ink-stained cells grown in capsule-induction medium as described in the Experimental Procedures. Scale bar = 5 μm. (B) Quantification of cell and capsule size following incubation in capsule-induction medium. Data represent the analysis for at least 60 cells for each strain with circles indicating individual data points and the black bar indicating the mean. (\*\*\**P* < 0.001). (C) 10-fold serial dilution assay of the indicated strains on L-DOPA agar plates to assess melanin production. Plates were incubated for 3 days at 30°C before being photographed. (D) The indicated strains were grown in liquid L-DOPA medium at 30°C for 24 h and melanin secretion was quantified by measuring the OD_405_ in the supernatant fraction. Data represent the mean ± SEM of three independent experiments. (\*\*\**P* < 0.001) (E) 10-fold serial dilution assay of the indicated strains on medium containing BFA (20 μg ml^−1^), monensin (1.25 mg ml^−1^), tunicamycin (0.15 μg ml^−1^), glycerol (2 %), ethanol (2 %), sodium acetate (1 or 2 %), H_2_O_2_ (1.5 mM), LiCl (100 mM), NaCl (1.5 M), KCl (1.5 M), congo red (2.5 mg ml^−1^) and calcofluor white (1 mg ml^−1^). All the plates were incubated for 3-4 days at 30°C or 37°C before being photographed.

### Additional components of clathrin-mediated endocytosis contribute to the use of heme and are required for virulence in mice

We also characterized the role of other components of CME in the use of iron from heme and hemoglobin. Strains deleted for the genes involved in different steps of CME were identified in the UCSF libraries and characterized for their ability to use different iron sources. Specifically, we tested mutants for the genes *LAS17, BBC1, YAP1801, SLA1, BZZ1, ARP2, CAP1, CAP2, SAC6, ARK1, PFY1, RVS161* and *RVS167*. A serial dilution spot assay on different iron sources revealed that strains lacking *LAS17* (encoding a homolog of Wiskott-Aldrich Syndrome protein; *WASP*) as well as *RVS161* and *RVS167* (encoding amphiphysin-like lipid raft proteins) showed impaired growth on low-iron media containing hemin as an iron source, similar to the *chc1*Δ mutant (Fig. 9A). Importantly, as observed for the *chc1*Δ mutant, the growth defects of the *las17*Δ, *rvs161*Δ and *rvs167*Δ mutants were also exclusively observed for the hemin and not for the inorganic iron sources, FeCl3 and FeSO4 (Fig. 9A). Notably, the inability to utilize iron from heme was even more pronounced in the *las17*Δ mutant compared to the *chc1*Δ strain, suggesting a crucial role of Las17 in heme-iron utilization in *C. neoformans*. To confirm this phenotype, we reconstructed two independent las17Δ mutants (*las17*Δ-*1* and *las17*Δ-*3*) in the H99 strain background and examined their growth on hemin or hemoglobin in liquid media assays. As expected, the *las17*Δ mutants were unable to grow properly in presence of either hemin or hemoglobin unlike the WT strain (Fig. 9B). These results further support a role for CME specifically in the use of iron from heme.

**Figure 9:**
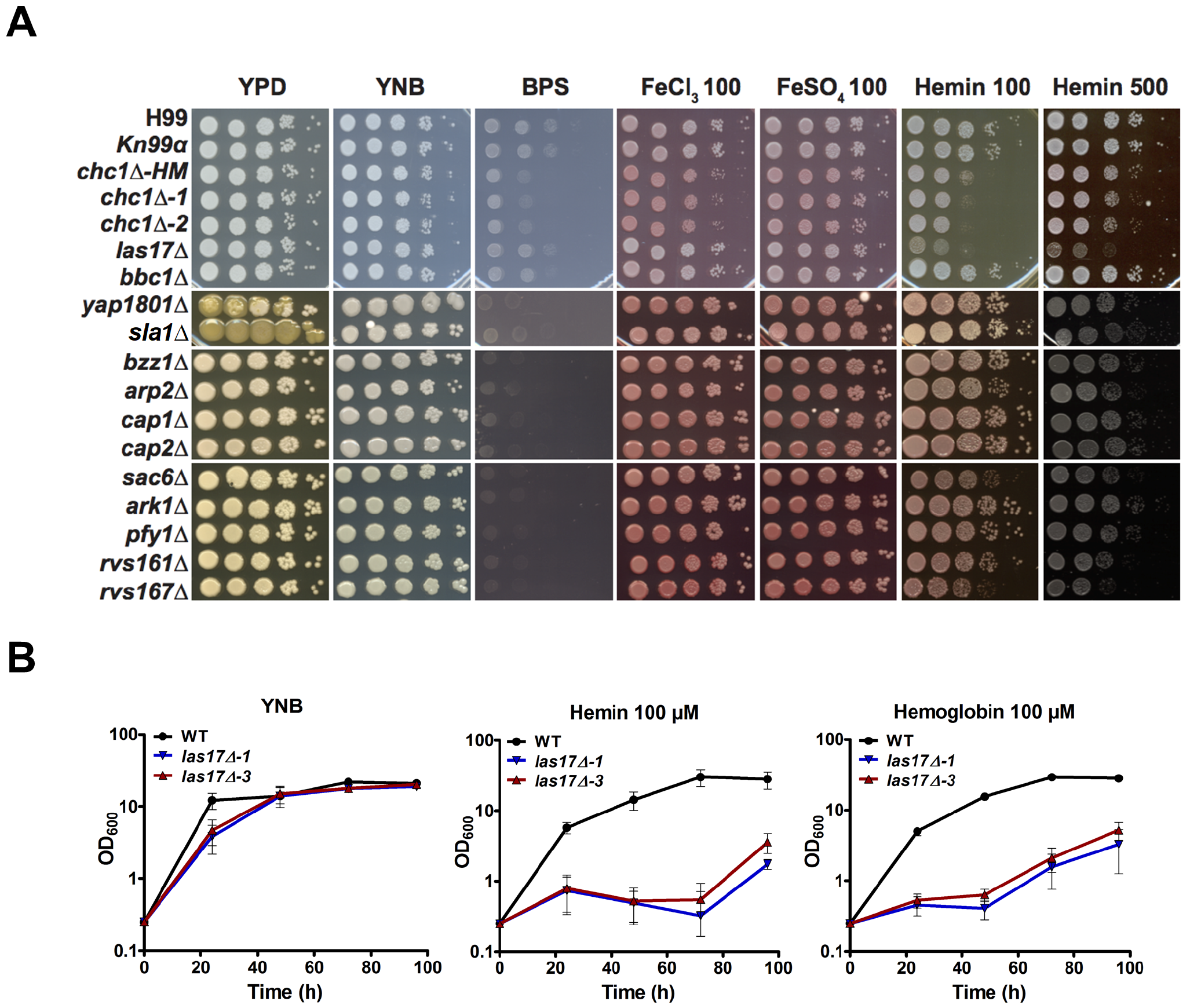
Las17, Rvs161 and Rvs167 are also required for heme-iron utilization in *C. neoformans*. **(A)** 10-fold serial dilutions of iron-starved cells of the indicated strains were spotted on medium supplemented as shown including BPS (100 μM), FeCl_3_ (100 μM), FeSO_4_ (100 μM) and hemin (100 or 500 μM). All of the plates were incubated for 3-4 days at 30°C before being photographed. (B) Iron-starved cells of the indicated strains were transferred to YNB-LIM medium containing either hemin or hemoglobin and their growth at 30°C was monitored by recording OD_600_ over a 96-h time-course at the indicated time intervals. Data are presented as the mean ± SEM of 3-4 independent experiments.

We focused on *las17*Δ for further phenotypic characterization. As observed for the *chc1*Δ mutant, the two independent *las17*Δ mutants also showed reduced growth in the presence of BFA and monensin and the salts NaCl, KCl and LiCl (Fig. 10A). Importantly, unlike the *chc1*Δ mutant, the *las17*Δ mutants did not show any growth inhibition at the host temperature of 37°C. Also, no growth defects were observed in utilizing alternate carbon sources, or in the presence of oxidative stress, ER stress and cell wall damaging agents. Finally, we also observed reduced capsule production and melanin secretion at 37°C in the *las17*Δ mutants (Figs. 10B and C). A previous study in the *C. neoformans* serotype D strain JEC21 revealed a role of *LAS17* (*CNBE5050*) in growth at 37°C, cytokinesis, chitin distribution, endocytosis and virulence (Shen *et al.*, 2011). We also conducted a virulence assay to compare the survival of mice infected with *las17*Δ mutants with the mice infected with the WT strain in H99 background. As shown in Figure 10D, mice infected with the *las17*Δ mutants showed significantly increased survival compared to the mice infected with the WT strain, although the mutants eventually caused disease. A preliminary test of the *rvs161*Δ and *rvs167*Δ mutants in three mice for each strain also indicated attenuated virulence (data not shown). We did not test the *chc1*Δ mutant in mice because of the growth defect at 37°C for this strain. Taken together, these results suggest that CME in *C. neoformans* not only is important in endocytosis but also plays essential roles in use of heme-iron, virulence and elaboration of several key virulence determinants including growth at high temperature, capsule and melanin production and survival in various stress conditions.

**Figure 10:**
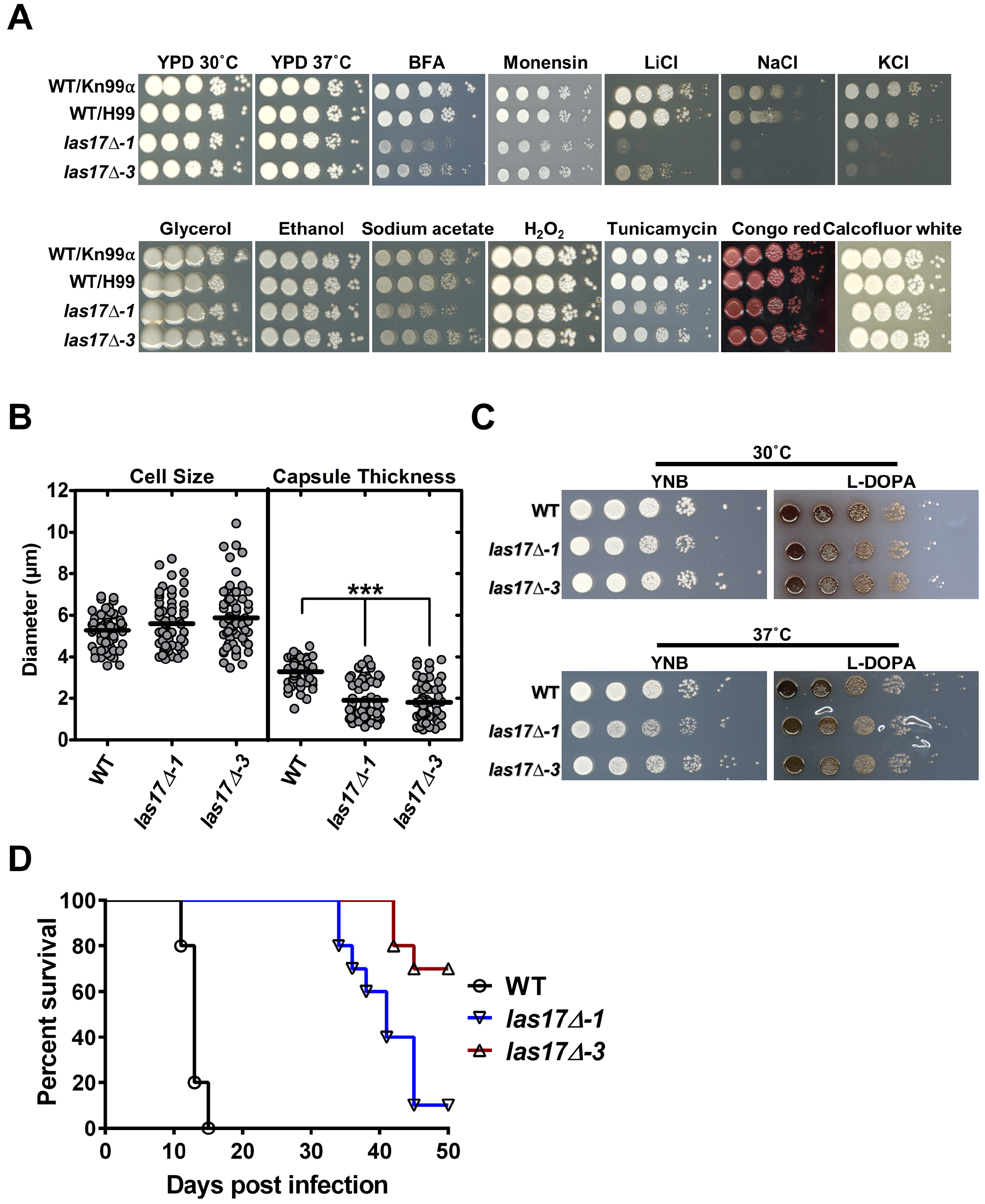
Clathrin-mediated endocytosis component Las17 is required for capsule and melanin production, and for virulence. (A) 10-fold serial dilutions of the strains were spotted on the indicated medium supplemented with different compounds as described in the legend of Figure 8E, with sodium acetate used at a concentration of 2%. All the plates were incubated for 3-4 days at 30°C or 37°C before being photographed. (B) Quantification of *C. neoformans* cell and capsule size following incubation in capsule-induction medium. Data represent the analysis of at least 60 cells for each strain with circles indicating individual data points and the black bar indicating the mean. (\*\*\**P* < 0.001). (C) 10-fold serial dilutions assay of the indicated strains on L-DOPA agar plates to assess the melanin production. Plates were incubated for 3 days at 30°C or 37°C before being photographed. (D) 10 female BALB/c mice were inoculated intranasally with 2 × 10^5^ cells of each of the strains indicated, and the survival of the mice was monitored daily. Survival differences between groups of mice were evaluated by log-rank tests. The **p**-values for the mice infected with the WT and mutant strains were statistically different (*p* < 0.001). Note that the experiment was terminated at 50 days when the mice infected with the mutants became ill.

## Discussion

Heme and hemoglobin serve as important iron sources for the proliferation of *C. neoformans* (Jung *et al.*, 2008; Bairwa *et al.*, 2017). However, very few factors have been identified for heme and hemoglobin uptake and internalization. Non-iron metalloprotoporphyrins are potential anti-microbial agents as well as useful reagents for characterizing heme uptake by microbes. In this study, we screened for mutants with altered susceptibility to manganese metalloporphyrin (MnMPP) and focused on the subsequent characterization of clathrin-mediated endocytosis because a *Chc1* mutant displayed enhanced growth on MnMPP and reduced growth on heme as an iron source. In addition to Chc1, we also identified other components of CME, including Las17, Rvs161 and Rvs167 that were necessary for efficient use of heme and hemoglobin as iron sources. Growth of the mutants lacking *CHC1* and *LAS17* in a medium with hemin or hemoglobin was strongly impaired, and growth was also significantly reduced for mutants lacking *RVS161* and *RVS167*. The *Chc1, las17, rvs161* and *rvs167* mutants also shared some of the conserved phenotypes seen for their counterparts in *S. cerevisiae*, *C. albicans* and *Aspergillus nidulans*, including defects in cell morphology, cell division, growth at high temperature and endocytic processes (Bauer *et al.*, 1993; Sivadon *et al.*, 1995; Naqvi *et al.*, 1998; Baggett and Wendland, 2001; Walther and Wendland, 2004; Douglas *et al.*, 2009; Schultzhaus *et al.*, 2017). Furthermore, lack of *CHC1, LAS17, RVS161* and *RVS167* in *C. neoformans* had a profound effect on key virulence factors viz. capsule and melanin production and, for *LAS17*, the ability to cause disease in a mouse model of cryptococcosis.

A role for CME in heme/hemoglobin utilization has been described in the intracellular parasite *Leishmania donovani* and the model worm *Caenorhabditis elegans* (Agarwal *et al.*, 2013; Yuan *et al.*, 2016). However, to date no role of CME in heme uptake/utilization has been reported for fungi. We showed that Chc1 in *C. neoformans* is found at the TGN and colocalizes with FM4-64 labelled vesicles, and this is consistent with observations in *S. cerevisiae* and mammalian cells (Puertollano *et al.*, 2003; Newpher *et al.*, 2005). Importantly, we observed the localization of Chc1 with intracellular hemoglobin-containing vesicles and this suggests that hemoglobin internalization in *C. neoformans* occurs via an endocytosis process involving clathrin-coated vesicle formation. Of course, there may also be CME-independent mechanisms of uptake. Furthermore, since CME occurs for receptor bound factors, our study highlights the possibility of cell-surface heme/hemoglobin binding factors in *C. neoformans*. In the pathogenic fungus *C. albicans*, proteins containing the CFEM (Common in Fungal Extracellular Membranes) domain including Rbt5, Rbt51, Csa2 and Pga7 play important roles in heme and hemoglobin binding, uptake and delivery into the cell (Weissman and Kornitzer, 2004; Okamoto-Shibayama *et al.*, 2014; Kuznets *et al.*, 2014; Nasser *et al.*, 2016). These cell surface-associated proteins function in a sophisticated relay network system to bind, extract and shuttle the heme molecule for its subsequent delivery into the cell (Kuznets *et al.*, 2014). In this regard, it is important to highlight that proteins containing the CFEM domain have been identified in *C. neoformans* (Zhang *et al.*, 2015). Whether or not these CFEM domain-containing proteins are involved in heme or hemoglobin cell surface binding and internalization in *C. neoformans* remains to be investigated.

Studies in both *C. albicans* and *C. neoformans* revealed that components of ESCRT-I, II, and III machinery are required for the use of heme as an iron source (Weissman *et al.*, 2008; Hu *et al.*, 2013; Hu *et al.*, 2015). Specifically, the ESCRT endocytic trafficking machinery proteins Vps23, Vps22 and Snf7 have been implicated in heme-iron use in *C. neoformans* (Hu *et al.*, 2013; Hu *et al.*, 2015). Based on our findings, we speculate that in *C. neoformans*, heme or hemoglobin is bound by as yet unknown cell surface-associated protein/s, followed by a coordinated assembly of clathrin-coated vesicles containing heme or hemoglobin-receptor that triggers sequential recruitment of Vps-proteins leading to endocytosis and trafficking of the heme/hemoglobin-receptor cargo to vacuole. A major step in internalization of cell-surface protein by CME involves mono-ubiquitination of cargo proteins (Hicke and Riezman, 1996; Egner and Kuchler, 1996; Goh *et al.*, 2010; Bertelsen *et al.*, 2011). Similarly, the ESCRT machinery mainly targets mono-ubiquitinated cell surface proteins for trafficking to the vacuole (Schuh and Audhya, 2014). Therefore, it is possible that a cell-surface protein involved in heme or hemoglobin binding in *C. neoformans* is also ubiquitinated and internalized by the CME and ESCRT pathways for its final delivery to vacuole. In this regard, it would be interesting to identify the structural components of clathrin-labelled hemoglobin-containing vesicles to potentially discover the predicted cell surface heme/hemoglobin receptor/s or transporter/s. Certainly, additional studies are required to delineate the molecular interactions between CME and ESCRT pathways to fully understand the intracellular trafficking of heme/hemoglobin. Unexpectedly, our metalloporphyrin screening did not reveal any potential transporters or cell-surface associated protein/s that could be directly involved in heme transport or binding. However, we did identify several candidate transport proteins including a multi-drug transporter, Pdr5, two uncharacterized drug efflux proteins encoded by *CNAG_03474* and *CNAG_01844*, a siderophore transporter, and sugar transporters. Further research is necessary to demonstrate any possible role of these transporters in heme-iron binding or transport, or in the efflux of toxic molecules such as MnMPP.

It is interesting that additional components of CME in *C. neoformans*, including the Las17 homolog of WASP, and the amphiphysins, Rvs161 and Rvs167, were also required for efficient use of heme/hemoglobin as iron sources. Individual mutants lacking these genes show significant growth defects in the presence of hemoglobin and this highlights the essential role of endocytosis in proper sorting and trafficking of heme cargo. We also observed defects in the expression of several key virulence factors in *chc1, las17, rvs161* and *rvs167* mutants including capsule elaboration and melanin secretion and attenuated virulence for the *las17* mutant in mice. In addition to heme-iron utilization, a contribution of CME to capsule elaboration is interesting because it suggests an important role in the trafficking of proteins and/or polysaccharides for the capsule synthesis. Similarly, impaired trafficking of laccase to the cell surface may account for the dependence of melanin formation on CME.

Environmental sensing of iron and heme availability is linked to intracellular signaling events leading to transcriptional and translational regulation of multiple factors involved in iron uptake, trafficking and metabolism, as well as the response to stress. Our metalloporphyrin screening also identified mutants deleted for the components of the Hog1 and Ca^2+^-calcineurin signaling networks including *hog1*Δ, *pbs2*Δ, *bem2*Δ, *sch9*Δ, *ste20*Δ, and *CNb1A*. Notably, both Ca^2^+-calcineurin and Hog1-mediated MAPK signaling in *C. neoformans* play important roles in the response to diverse environmental stresses (e.g., osmotic, oxidative, antifungal drug, heavy metal, and high temperature stress), and in the production of capsule and melanin (Kozubowski *et al.*, 2009; Jung and Bahn, 2009; Kronstad *et al.*, 2013). Additionally, the Hog1-dependent MAPK signaling pathway is involved in iron uptake and homeostasis in *S. cerevisiae* and in the fungal pathogens *C. albicans, C. glabrata* and *A. fumigatus* (Kaba *et al.*, 2013; Srivastava Vivek K. *et al.*, 2015; Castro *et al.*, 2016; Martins *et al.*, 2018). It is plausible that a deficiency for iron or heme-iron in *C. neoformans* also acts as an external/internal stress signal that induces activation of major signaling events, and this may eventually result in modification of the iron-uptake machinery or intracellular iron metabolism. However, the functions of different components of the Hog1 and calcineurin signaling pathways in heme-iron uptake and metabolism and their potential connection with the clathrin-mediated endocytosis and heme-iron trafficking remain to be elucidated.

In conclusion, our study highlights the value of genome-wide screening approaches in a human fungal pathogen for not only identifying new contributions of CME and signaling pathways in heme-iron utilization, but also potential applications of non-iron metalloporphyrins in the treatment of cryptococcosis. Importantly, the contributions of CME and MAPK/calcineurin-signaling for heme/hemoglobin uptake and metabolism in *C. neoformans* appear to be evolutionary conserved with yeast and higher eukaryotes. Further exploration of the molecular mechanisms for heme sensing, uptake, endocytosis, signaling and regulation will provide a deeper understanding of iron metabolism in fungal pathogens and subsequently assist in the development of new therapeutic targets.

## Experimental Procedures

### Strains, plasmids, media and growth conditions

*C. neoformans* var. *grubii* strains H99 (serotype A) and Kn99-alpha were used as WT parental strains. *C. neoformans* strains were routinely maintained on YPD agar medium (1% yeast extract, 2% Bacto-peptone, 2% D-glucose and 2% agar). Overnight cultures were obtained by inoculating a single colony into liquid YPD medium and growing at 30°C with shaking at 200 revolutions per min (rpm). YPD medium plates containing nourseothricin (*NAT*) (200 μg ml^−1^) were used to select the transformants deleted for the *CHC1* and *LAS17* genes. Solid YPD medium containing neomycin (*NEO*) was used to select the *chc1*Δ/*CHC1-GFP* complemented strain. The neomycin resistance cassette and the GFP gene was from the plasmid pWH091 as described earlier (Jung *et al.*, 2009). Low iron YNB medium (Yeast Nitrogen Base with amino acids; YNB-LIM) was prepared by adding the iron-chelator bathophenanthroline disulfonate (BPS; 150 μM) in YNB medium prepared in Chelex 100 resin-treated low iron water. All chemicals were obtained from Sigma-Aldrich (St. Louis, MO) unless indicated otherwise. The strains used in this study are listed in Supplementary Information Table S2.

### Construction of deletion mutants and complemented strains

The candidate genes *CHC1* (clathrin heavy chain) and *LAS17* (Wiskott-Aldrich syndrome protein (WASP) homolog) were deleted in the *C. neoformans* strain H99 by homologous recombination using gene-specific knockout cassettes containing the resistance marker nourseothricin and the 5’ and 3’ flanking sequences of the respective genes. These gene-specific knockout cassettes were amplified from genomic DNA of *chc1*Δ and *las17*Δ mutants in Kn99α background (mutants obtained in UCSF-libraries) using primers GB-04904-ForKO, GB-04904-RevKO and GB-02029-ForKO, GB-02029-RevKO, respectively. The constructs were introduced into the *C. neoformans* H99 wild type strain by biolistic transformation, as described previously (Davidson *et al.*, 2000). Two independent *chc1*Δ mutants (*chc1*Δ-*1* and *chc1*Δ-*2*) and two *las17*Δ mutants (*las17*Δ-*1* and *las17*Δ-*3*) were selected and evaluated in the experiments. Complementation of the *chc1*Δ mutation was accomplished with a modified overlapping PCR strategy to include the green fluorescence protein sequence (GFP) at the C-terminus of Chc1 before the stop codon. Briefly, the left arm (~1 kb amplicon of the 3’ end of *CHC1* ending just before stop codon; amplicon-1) and right arm (~1 kb amplicon starting after the stop codon of *CHC1*; amplicon-2) from the *CHC1* genomic locus were amplified from WT genomic DNA using the primers GB-04904-For and GB-04904-GFP-Rev (containing an 18 base overlap with the GFP sequence at the 5’ end) and GB-04904-3′UTR-GFP-For (containing a 20 base overlap with the GFP sequence at the 3’ end) and GB-04904-3′UTR-Rev, respectively. The GFP gene and the neomycin resistance cassette (amplicon-3) were amplified from the plasmid pGH025 (Hu *et al.*, 2017) using primers GB-CnGFP-Neo-For and GB-CnGFP-Neo-Rev. Overlap PCR was performed using amplicon-1, −2 and −3 as templates with primers GB-04904-For and GB-04904-3′UTR-Rev to obtain the *CHC1-GFP-NEO-3’UTR* fusion product. This fusion product was transformed into the WT strain by biolistic transformation and the positive transformants were selected for resistance to G418 (Geneticin, Sigma).

For complementation of *chc1*Δ with the *CHC1-GFP* sequence, the *CHC1* genomic locus from WT DNA containing the *CHC1-GFP* variant was amplified with primers GB_04904_For_Comp and GB_04904_Rev_Comp and transformed into the *chc1*Δ strain. The positive transformants were selected for resistance to G418. Proper location and orientation of the gene fusions at the *CHC1* locus were determined by PCR. Primers used in this study are listed in Supplementary Information Table S3.

### Metalloporphyrin screening and identification of mutants with altered MnMPP susceptibility

To identify new factors involved in heme-iron use and susceptibility to manganese metalloporphyrin, we designed the screening strategy outlined in Figure 1B. Two libraries of deletion strains of *C. neoformans* (~3,300 genes; arranged in 96-well plates) constructed by Dr. Hiten Madhani’s group at the University of California, San Francisco, USA, were obtained from the Fungal Genetics Stock Center (http://www.fgsc.net/crypto/crypto.htm). These libraries include a set of 14 plates of deletion strains submitted to FGSC in year 2008 (UCSF-2008) and another set of 22 plates of deletion strains submitted in 2015 (UCSF-2015). Additionally, a library of ~322 signature-tagged gene deletion strains for 155 putative transcription factors (TF) of *C. neoformans* (at least two independent deletion strains per TF; TFKO) was constructed by Dr. Yong-Sun Bahn’s group at Yonsei University, Korea and obtained from Dr. Won Hee Jung (Chung-Ang University, Korea). In this study, these three libraries (UCSF-2008, UCSF-2015 and TFKO) were screened for susceptibility to manganese metalloporphyrin (MnMPP). To initiate the screening, deletion mutants were grown in 96-well plates in YPD at 30°C overnight. These YPD precultures were iron-starved in 96-well plates containing low-iron YNB medium (YNB with BPS-150 μM) for two rounds of one day each. After iron starvation, 10 μl of culture was introduced into the test media conditions including regular YNB, YNB with BPS (150 μM), YNB with BPS and hemin (10 μM) [Hemin-medium] and YNB with BPS supplemented with hemin (10 μM) and MnMPP (10 μM) [MnMPP-medium] in a final volume of 200 μl. The plates were incubated at 30°C with shaking at 50 rpm and the OD_600_ was measured after 72 h using a microplate reader (Infinite M200; Tecan). YNB and YNB with BPS (150 μM) were used as control plates to evaluate the growth of the strains with and without iron starvation. Any strain that failed to grow in YNB was removed from the analysis.

*C. neoformans* deletion mutants showing differential susceptibility to MnMPP were identified by assessing the 72 h growth differences between the hemin with MnMPP medium versus hemin only medium conditions. The OD600 values in these two media conditions were normalized with their respective blank media controls and the relative growth difference (in percentage) for each strain was calculated using the equation: ((OD_600_ of the strain in MnMPP medium at 72 h/OD_600_ of the strain in hemin medium at 72 h) × 100). An average of all the percentage relative growth difference values from all the strains in each library was determined (Average Relative Growth; ARG, ARG^4^ for 4 plates in TFKO library, ARG^14^ for UCSF-2008 library, ARG^22^ for UCSF-2015 library) to evaluate the overall growth behavior of the strains in each individual library. To identify the strains with altered MnMPP susceptibility, we used a cut-off of 1.5-fold difference between the individual percentage growth and the ARG value of the respective library. The strains that grew ≥ 150% or ≤ 50% of the ARG values were selected as MnMPP resistant and sensitive strains, respectively.

### Gene Ontology enrichment and STRING network analysis

Gene Ontology (GO) analysis for the genes identified in the screens was performed using the PANTHER gene ontology online tool (http://pantherdb.org). The genes were evaluated for enrichment of GO biological processes, molecular functions and cellular component categories by comparing with a background genome set of *C. neoformans* present in the PANTHER database with a cut-off of *P*<0.05. The interaction network for the genes was obtained from the STRING (https://string-db.org) online network analysis with the *C. neoformans* var. *neoformans* JEC21 genome database as a background using all active interaction sources and a medium confidence value of 0.400. The network that was obtained was extracted and visualized using the Cytoscape network analysis tool (http://www.cytoscape.org). The cluster identification in Cytoscape was done using the Community Cluster (GLay) network clustering algorithm in the clusterMaker plugin.

### Serial dilution spotting and liquid growth assays

The ability of mutants to grow under different environmental conditions was evaluated by the spotting of serial dilutions on solid agar plates and by optical density (OD_600_) measurement in liquid growth assays. For serial dilution spotting assays, the cells were grown in YPD at 30°C and 200 rpm for 14-16 h, washed three times with either PBS or Chelex 100 resin-treated low iron water and adjusted to 2 × 10^7^ cells ml^−1^. The cell suspension was diluted 10-fold serially and 5 μl of each dilution was spotted onto YPD and/or YNB plates supplemented with different compounds. Plates were incubated at 30°C or 37°C for 2-5 days before being photographed. Strains were evaluated for growth responses toward temperature (30°C and 37°C), salts (1.5 M NaCl, 1.5 M KCl, 100 mM LiCl), trafficking inhibitors (20 μg ml^−1^ brefeldin A; BFA and 1.25 mg ml^−1^ monensin), alternate carbon sources (2% of either glycerol, ethanol or sodium acetate), antifungal drugs (5, 7.5 or 10 μg ml^−1^ fluconazole) and agents that cause the unfolded protein response in the endoplasmic reticulum (0.15 μg ml^−1^ tunicamycin and 10 mM dithiothreitol; DTT), alter redox state (1 or 1.5 mM H_2_O_2_) and challenge cell wall integrity (1 mg ml^−1^ calcofluor white and 2.5 mg ml^−1^ congo red).

To evaluate the growth on different iron sources, cells were starved for iron before growth evaluation, as described earlier with slight modifications (Hu *et al.*, 2017). Briefly, overnight YPD-grown fungal cells were washed three times in Chelex 100 resin-treated low iron water and inoculated into OD_600_ of 0.2 in YNB with BPS (150 μM, YNB-LIM). The cells were incubated for 24 h before spotting onto YNB and YNB-LIM solid agar plates with and without supplemented inorganic iron sources (10 or 100 μM of FeCl_3_ or FeSO_4_) or hemin (10, 100 or 1000 μM). Liquid growth assays were performed in conical flasks with YNB and YNB-LIM media with and without supplementation of hemin and hemoglobin at different concentrations. Iron-starved cells (one day) were inoculated to OD_600_ of 0.2 in fresh media. The total medium volume was 20 ml. The flasks were incubated in a shaker at 30°C and 200 rpm and OD_600_ was measured at regular time intervals.

### Capsule formation and melanin production

To evaluate capsule formation, cells were grown overnight in YPD, washed with water and incubated in low-iron capsule induction medium (CIM; 5 g liter^−1^ glucose, 5 g liter^−1^ L-asparagine, 0.4 g liter^−1^ K_2_HPO_4_, 0.25 g liter^−1^ CaCl_2_.2H_2_O, 0.08 g liter^−1^ MgSO_4_.7H_2_O, 4.78 g liter^−1^ HEPES, 1.85 g liter^−1^ NaHCO_3_ [dissolved in Chelex 100 resin-treated water], pH 7.4) at 30°C and 200 rpm for 48 h. To visualize the capsule, cells were stained with India ink and observed by differential interference contrast (DIC) microscopy (Zeiss Axioplan 2) at 100X resolution. Capsule and cell sizes were measured in ZEN lite software (ZEISS Inc.) from at least 50 to 60 cells for each strain.

Melanin production was evaluated on either solid or liquid L-3,4-dihydroxyphenylalanine (L-DOPA; Sigma) media (1 g liter^−1^ L-asparagine, 3 g liter^−1^ KH_2_PO_4_, 0.25 g liter^−1^ MgSO_4_.7H_2_O, 0.2 g liter^−1^ L-DOPA and 0.0001% thiamine hydrochloride [dissolved in distilled water], pH 5.6) containing 0.1% glucose. For melanin production on solid media, serial dilution spotting of the overnight YPD-grown cells was done on L-DOPA solid agar plates as described above. The plates were incubated in the dark at 30°C or 37°C for 48 h before being photographed. Melanin secretion in the liquid L-DOPA medium was analyzed as described earlier with some modifications (Santiago-Tirado *et al.*, 2015). Briefly, YPD-grown overnight cells were collected, washed twice in Chelex 100 resin-treated low-iron water and adjusted to 10^7^ cells ml^−1^ in low-iron water. 300 μl of the cell suspension was added to 3 ml L-DOPA and samples were incubated in the dark at 30°C and 200 rpm for 24 h. After incubation, 1 ml culture was centrifuged at 1,000 g for 5 min and the OD_405_ was measured for the supernatant fractions to analyze the melanin secretion.

### FM4-64 and Solophenyl Flavine (SF) labelling and microscopy

To examine the endocytosis process, trafficking of the lipophilic styryl dye, N-(3-triethylammoniumpropyl)-4-(6-(4-(diethylamino) phenyl-hexatrienyl) pyridinium dibromide (FM 4-64, Molecular Probes) was examined as described earlier with some modifications (Vida and Emr, 1995). Briefly, YPD-grown overnight cells were collected, washed twice with cold-PBS and incubated in cold-PBS with FM4-64 (20 μM) at 4°C for 30 min. The cells were harvested at 1,000 g for 3 min at 4°C, washed three times with ice cold-PBS, resuspended in fresh YPD medium and further incubated at RT. At different time intervals, a small portion of the cells was withdrawn from the YPD suspension and placed on a standard glass slide for visualization under a fluorescence microscopy (Zeiss Axioplan 2) with rhodamine filter.

To visualize chitin, exponentially growing cells were labelled with solophenyl flavine (SF) in PBS for 10 min at room temperature as described earlier (Hoch *et al.*, 2005). Labelled cells were visualized under a fluorescence microscopy using the EGFP filter set. All photomicrographs were captured at 100X resolution using a Plan DIC achromatic objective and analyzed with ZEN lite software.

### Labelling of hemoglobin with Alexa Fluor 594 and analysis of the kinetics of intracellular trafficking

The uptake and intracellular trafficking of hemoglobin (Hb) in *C. neoformans* cells was examined with labelled hemoglobin. Human hemoglobin (Sigma) was labelled with Alexa Fluor 594 according to the manufacturer’s protocol (Invitrogen, Molecular Probes). Next, exponentially growing cells were collected, washed twice with ice-cold PBS and incubated in PBS containing Alexa Fluor 594-labelled hemoglobin (Hb^Alexa594^, 50 μg/ml) for 20 min on ice to allow cell surface binding. After 20 min, cells were washed three times with ice-cold PBS to remove unbound Hb^Alexa594^ and resuspended in PBS prewarmed at 30°C. After incubation at different time points, cells were visualized under a fluorescence microscope (Zeiss Axioplan 2) using Alexa Fluor 594 filter set with exposure time of 5000-10000 milliseconds. For Chc1-GFP and Hb^Alexa594^ co-localization experiments, cells expressing Chc1-GFP were labelled with Hb^Alexa594^ as described and visualized with Alexa Fluor 594 and EGFP filter sets at exposure times of 5000 and 1000 milliseconds, respectively. All photomicrographs were captured at 100X resolution using a Plan DIC achromatic objective and analyzed with ZEN lite software.

### Total cellular heme level quantification

Total heme content of the cells was determined using the BioVision Hemin assay kit (BioVision, Milpitas, California) as described earlier (Do *et al.*, 2017). Briefly, cells were grown in YPD medium overnight at 30°C, washed twice with PBS and lysed using bead beating in buffer containing Tris-HCl (10 mM, pH 8.0) and NaCl (150 mM). After lysis, supernatant was collected by centrifugation at 13,000 rpm for 3 min at 4°C and used to measure the heme content as described by the manufacturer. Total heme level in each strain was normalized to the protein concentration.

### Virulence assay

A mouse inhalation model of cryptococcosis was used to assess the virulence of mutants using 4-6 weeks old female BALB/c mice, as described earlier (Hu *et al.*, 2017). Briefly, strains were grown overnight in YPD medium at 30°C, washed and resuspended in PBS. A total of 2 × 10^5^ cells in a 50 μl volume of PBS were used for intranasal inoculation of 10 mice per strain. The status of each mouse was monitored daily post-inoculation. Mice reaching the humane endpoint were euthanized by CO_2_ anoxia. The protocol for the virulence assay (protocol A17-0117) was approved by the University of British Columbia Committee on Animal Care.

### Statistics

Data were visualized and statistically analyzed using GraphPad Prism version 6.0 (Graph-Pad Software, Inc., USA). Statistical tests were performed by one-way analysis of variance (ANOVA) followed by a Bonferroni correction for melanin secretion assays, or by Student’s t test for capsule size, or by log-rank tests for *in vivo* mouse survival assay. *P* values of 0.05 were considered to be significant.

## Acknowledgements

This research was supported by grant 5R01AI053721 from the National Institute of Allergy and Infectious Disease (to J.W.K.). J.W.K. is a Burroughs Wellcome Fund Scholar in Molecular Pathogenic Mycology. L.H. is the recipient of a doctoral scholarship from the National Sciences and Engineering Research Council of Canada. We thank the groups of Drs. H. Madhani and Y.S. Bahn for the construction and distribution of deletion mutant collections, and Dr. François Mayer for helpful comments on the manuscript.

## Author’s contribution

G.B. and J.W.K. conceived the study and designed the experiments. G.B. performed the experiments. G.B. and J.W.K. analysed the data and wrote the manuscript. G.H., M.C. and L.H. conducted the virulence assays in mice and contributed to the preparation of the manuscript.

## Supporting Information Figures legends

**Supporting Information Figure S1. *C. neoformans* MnMPP resistant mutants show susceptibility to diverse environmental growth conditions.** Heatmap depicting the growth behavior of selected MnMPP resistant mutants against various environmental stress conditions. The growth of the individual strains at 30°C, 37°C and in presence of hemin (10 μM), NaCl (1.5 M), brefeldin A; BFA (20 μg ml^−1^), tunicamycin (0.15 μg ml^−1^), dithiothreitol; DTT (10 mM), H_2_O_2_ (1 mM), calcofluor white (1 mg ml^−1^), and congo red (2.5 mg ml^−1^) was checked and displayed with color gradients.

**Supporting Information Figure S2. The *C. neoformans chc1*Δ mutant exhibits growth sensitivity to the iron-chelating drug curcumin.** 10-fold serial dilutions of iron-starved cells of the indicated strains were plated on medium supplemented as shown; BPS (100 μM), curcumin (300 μM), hemin (100 μM), FeCl_3_ (100 μM) and FeSO_4_ (100 μM). All the plates were incubated for 2-3 days at 30°C before being photographed.

**Supporting Information Figure S3. Trafficking inhibitor brefeldin A (BFA) affects Chc1 localization.** (A) Exponentially growing cells of wild-type stain expressing Chc1-GFP were treated with BFA (20 μg ml^−1^) for 1 h and localization of Chc1-GFP was visualized at the indicated time intervals after the treatment. Scale bar = 2 μm. (B) Quantification of the cells showing Chc1-GFP punctate structures on cell surface. 30-50 cells were analyzed from two experiments. Data represent mean ± standard deviation (SD) of the two experiments. (C) Chc1-GFP visualization in different media conditions. Strain expressing Chc1-GFP was incubated in the indicated media for 1 h before visualization under a fluorescence microscope. Scale bar = 2 μm.

